# Modulation of repopulating microglia in multiple sclerosis models with implication for neuroprotection

**DOI:** 10.1101/2025.08.08.669252

**Authors:** Neele Heitmann, Sarina Fiene, Lina Rambuscheck, Britta Eggers, Maria Fernanda Valdes Michel, Anne-Christin Gude, Hasan Hüsein Hendek, Sarah-Marie Oberhagemann, Katharina Klöster, Robert Hoffrogge, Dorsa Moezzi, Anna Karachunskaya, Hannah Märte, Kamilla Adam, Svitlana Rozanova, Verian Bader, Carlos Plaza-Sirvent, Gayatri Tandon, Martin Eisenacher, Jason R. Plemel, Konstanze F. Winklhofer, Oliver Gramlich, Katrin Marcus-Alic, Ingo Schmitz, V. Wee Yong, Ralf Gold, Simon Faissner

## Abstract

Microglia play a critical role in CNS pathologies, including multiple sclerosis (MS), and their modulation offers therapeutic potential, especially during progressive disease courses.

Using cell culture and experimental autoimmune encephalomyelitis (EAE) models, we investigated microglial dynamics during depletion and repopulation and their modulation using siponimod, an established CNS-penetrating MS medication.

Repopulating microglia exhibited a transient reactive state (CD86, MHC-II, *Il1b, Tnf)*. Siponimod modulated microglia populations, increasing CD163^+^ and CX3CR1^+^ while reducing CD86^+^MHC-II^+^ cells, accompanied by a reduction of neuronal damage. Proteomic spinal cord analysis revealed protein expression alterations by microglia repopulation and siponimod linked to inflammation, myelination, and neuronal structural organization, supported by RNA sequencing of the spinal cord. The neuroinflammation-attenuating role of siponimod could be linked to cell maintenance and myelin formation-associated processes.

These findings highlight the capacity of pharmacological interventions to modulate microglia, offering new insights into therapeutic strategies targeting microglial activity in neuroinflammatory diseases.

## Introduction

Multiple sclerosis (MS) is a neuroinflammatory demyelinating disease of the CNS (1), characterized by demyelinating lesions in the brain, spinal cord, and optic nerve. The disease is modulated by autoreactive T and B lymphocytes and influx of macrophages with consecutive neurodegeneration (2). Immune cells within the lesions interact with microglia, promoting a more inflammatory microglia polarization (3, 1). Most patients initially present with a relapsing remitting MS phenotype (RRMS) (4). Without treatment, RRMS evolves to a progressive disease course with fewer relapses but continuous disability accumulation, secondary progressive MS (SPMS) (4). In a smaller number of people, the disease progresses without relapse activity as primary progressive MS (PPMS). The understanding and treatment of MS have evolved greatly over the past decades, enabling the implementation of different medications, especially for RRMS, and improving the quality of life and the life expectancy of people living with MS (pwMS) (4, 5). While treatment of RRMS has made tremendous headway during the last 20 years, treatment of progressive MS is still unsatisfactory (5, 6) with only limited options, including B cell depletion (ocrelizumab, PPMS), siponimod (sipo, SPMS) (4) and the 2026 expected BTK-inhibitor tolebrutinib (7). This emphasizes the present necessity to enhance comprehension of progression in MS, particularly in isolation from relapse activity, with a view to facilitating the development of more efficacious drugs to mitigate progression.

CNS-based inflammation is a feature of progressive MS, involving microglia, astrocytes and B- and T-cell activation (8, 4). Microglial participation in inflammatory responses is characterized by cytokine release and oxidative stress. However, these cells also play a crucial role in maintaining normal brain function and facilitating regeneration following damage (4). Analysis of human microglia in pwMS is limited to biopsies, sophisticated imaging methods, and *post-mortem* tissue, and shows high microglial prevalence both within lesions and in normal-appearing white and grey matter during disease progression. Microglia at lesion edges can take up iron, associated with the enlargement of iron rim lesions, which are associated with faster progression. Moreover, microglia metabolize lipids, leading to the formation of foamy microglia in response to myelin debris (4, 9).

In addition, leptomeningeal inflammation and meningeal B-cell aggregates can induce cortical microglial activation and neurodegeneration, further supporting a role for microglia as key effectors within a complex inflammatory milieu. However, these data also underscore that progressive MS pathogenesis involves multiple interacting components, including B and T cells, astrocytes and other CNS-resident and infiltrating immune cells, and that the mechanisms shaping inflammation in progressive MS remain only partially understood.

In addition to the restriction of lymphocyte infiltration, microglia modulation is a promising treatment strategy that is currently being explored for the development of new therapeutics, including BTK inhibitors. Drugs that modulate microglia and other cells in the CNS must be able to cross the blood-brain barrier. Sipo, a medication authorized for the treatment of SPMS, exerts effects on CNS immune cells. Sipo mediates functional antagonism at the sphingosine-1-phosphate receptor (S1PR) 1, preventing immune cell egress from lymphoid organs and limiting CNS infiltration (10). Sipo has the ability to cross the blood-brain-barrier and modulates S1PR1 as well as S1PR5, both of which are present on different cell types in the CNS, including microglia (11). These properties promise its ability to modify or limit progression (4). The potential beneficial properties of sipo have been investigated in previous studies, but have varied in species (12, 11, 13), route of administration (14), and analysis.

The objective of this study was to determine the microglial phenotype during repopulation in a model of MS, experimental autoimmune encephalomyelitis (EAE), and to assess whether sipo could regulate microglia to establish a neuroprotective environment, with implications for disease progression. Given that sipo is already approved for all forms of clinically active MS, the detection of a new mode of action has the potential to provide insights into the long-term effects of the drug within the CNS that may be applicable to other diseases with microglial involvement, including neurodegenerative disorders.

## Results

The impact of sipo on CNS cells in pwMS remains unconfirmed. While sipo is an effective treatment for pwMS by reducing the leukocyte infiltration into the CNS, previous data suggest additional protective properties when reaching the CNS in a suitable concentration to agonize the S1PRs on CNS resident cells (14, 12). We examined sipo’s protective potential against stressors *in vitro*, without exerting adverse effects.

### Sipo prevents iron toxicity in human and murine cell cultures

Sipo was tested in human and murine cells. The treatment and stimulation were applied similarly to both species (Figure 1a,h). In live-dead cell imaging, sipo did not increase the percentage of dead cells of neurons and in neuron-microglia co-cultures (Figure S1a,c,e,h). The fold change in dead cells treated with sipo did not exceed that of the control group, both in the human and the murine system over time and at a late time point (Figure S1b,d,f,g,I,j). As iron deposition activates microglia in iron rim lesions (15), iron was used as a stressor to simulate the MS-diseased brain. Neurons were pretreated with sipo before the addition of microglia and subsequently iron. In human neuron cultures, iron induced 16.6% more dead cells after 24 hours, which was reduced by 10.5% with 1 µM sipo pretreatment (Figure 1b-d). In human neuron-microglia co-cultures, iron-induced cell death increased at about 16 hours after initiation of iron stimulation (Figure 1e,f). After 20 hours, iron induced roughly a two-fold increase in dead cells (Figure 1g, p<0.0001). Pretreatment with sipo protected against iron toxicity (p<0.0001; Figure 1g).

**Figure 1.**
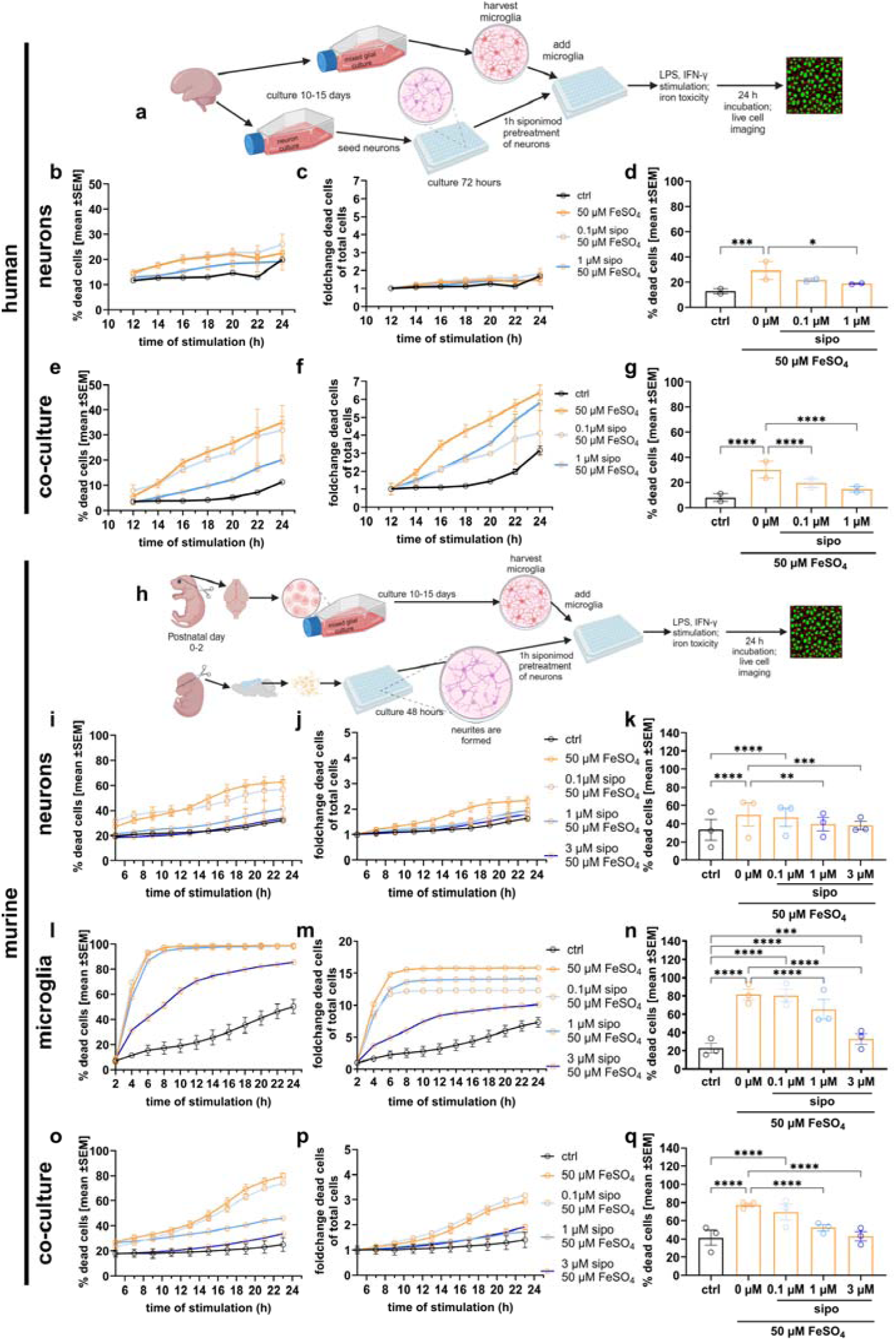
Iron toxicity studies in cell culture. Primary human and murine neuron and microglia cell cultures (a,h) stimulated with FeSO_4_ and treated with siponimod (sipo) were imaged with live-dead-staining every two hours (b,c human neurons; e,f human co-culture; I,j murine neurons; l,m murine microglia; o,p murine co-culture). Statistical analysis of specific time points in the human culture are presented after 24 hours (d) and 20 hours (g) of stimulation of two independent experiments. In murine cultures, the analysis was conducted with the data from time points after stimulation (k 22 h,n 6 h,q 22h) with three independent experiments. Experiments were measured in triplicates and analyzed with mean, SEM and n per experiment with the 2-way ANOVA and Šídák’s multiple comparisons test. Co-culture – neuron and microglia culture, ctrl – control. a and h were created with Biorender. P-values are depicted as * - p≤0.05, ** - p≤0.01, *** - p≤0.001, **** - p≤0.0001.

In the murine system, neuron, microglia, and neuron-microglia co-cultures were tested for susceptibility to iron toxicity. Iron induced cell death in all cultures (Figure 1i,l,o). As in human cultures, cell death was most pronounced in microglia and microglia-neuronal co-cultures with no effect of 0.1 µM sipo (Figure 1j,k,l,n,p,q). Both 1 and 3 µM sipo significantly reduced cell death after 22 hours of iron stimulation (Figure 1n,q).

The neuroprotective effects of sipo against iron toxicity in the co-culture were evaluated by analysis of neuronal morphology (Figure S2a). One µM and 3 µM sipo treatment ameliorated the reduced fiber length by 6-6.5% but failed to reach statistical significance (Figure S2b). Immunostaining of microglia revealed that iron reduced the proportion of microglia, increased the percentage of iNOS^+^ microglia, and reduced ramification (Figure S2c-d). With 1 and 3 µM sipo pretreatment, these iron impacts on microglia were reversed.

Upon stimulation of microglia with IFN-γ and LPS (iLPS), an increase of iNOS-expressing microglia in culture was observed along with their increased size, with no adverse effects on neuron or microglia health and minor reversing effects by sipo (Figure S2f-I, Figure S1k-p). To assess the effects of sipo on IFN-γ stimulation, we performed flow cytometry and observed a reduction in CD86 expression by sipo (Figure S2j). Given that sipo exerts greater effects on microglia than on neurons, we tested whether sipo could also prevent microglial stimulation in microglia-splenocyte co-cultures, which mimic the infiltration of peripheral immune cells into the CNS and microglial stimulation (Figure S2k). Although there were more antigen-presenting lymphocytes upon PMA/Iono stimulation, microglia populations were not differentially affected (Figure S2l,m). Sipo pretreatment reduced the frequencies of CD86^+^ microglia and significantly elevated the homeostatic marker CX3CR1 (Figure S2n). In summary, sipo demonstrated protective effects against iron-induced cell death in both human and murine neuronal and microglial cultures, as well as against inflammatory stimuli such as iLPS and splenocytes, reducing cell death and modulating microglial activation, morphology, and function.

### Low-dose sipo does not modulate brain microglia in EAE

The objective of microglia depletion during EAE was to deplete disease-mediating microglia, thereby creating conditions conducive to subsequent microglia repopulation. The tissue explant was conducted five days after the last microglia-ablating DT injection, when the microglia repopulation (MG^repo^) started. The hypothesis was that repopulating microglia would not become disease-promoting and would be susceptible to modulation by sipo. A subclinical dosage of sipo was administered to achieve sufficient concentrations in the CNS while allowing the investigation of whether MG^repo^ would modulate EAE (Figure S4a). To evaluate microglia depletion during EAE and to exclude adverse effects of tamoxifen or DTR expression (16), control experiments were performed (Figure S3a), confirming no adverse effects and identifying post-remission as the most tolerable timing for MG^repo^ (Figure S3b-f). MG^repo^ was confirmed by flow cytometry of brain tissue, which documented repopulation of CD86^+^ microglia (Figure S3g,h). 0.45 µg sipo reduced EAE incidence and severity but did not prevent EAE-induced weight loss (Figure S4b,c,d). In MG^repo^ vehicle and MG^repo^ sipo, microglia, macrophages, and lymphocytes displayed higher CD86^+^ cell frequencies in the brain (Figure S4e). Although the cell culture data suggested that sipo modulates microglia, this effect was not demonstrated at a sipo dose of 0.45 µg per day during EAE. Furthermore, the substantial prevention of lymphocyte egress from lymphoid organs was not achieved. (Figure S4f). In summary, low-dose sipo had a moderate effect on EAE disease severity and peripheral immune cells and failed to prevent the presence of CD86^+^ microglia in the brain.

### Sipo ameliorates EAE disease course and spinal cord pathology by neuroprotection

A higher dose of 1 µg sipo was administered in EAE to enable higher concentrations to reach the CNS and exert direct effects on CNS-resident cells, such as microglia (Figure 2a). Here, sipo reduced the EAE incidence (Figure 2b) and ameliorated overall EAE severity by preventing weight loss, ameliorating EAE scores, and reducing the peak of disease (Figure 2c-e). MG^repo^ led to cerebrospinal ataxia three days after the last DT injection and remained unaffected by sipo (Figure 2f). The spinal cord pathology was assessed initially by combining H&E with LFB staining (Figure S5a-c). The area of infiltrating leukocytes (H&E) was significantly reduced by sipo, and MG^repo^ was reflected by CD3^+^ T cells and GFAP intensity-based astrogliosis (Figure S5d, Figure 2g,h). Flow cytometry revealed that sipo limited lymphocyte and macrophage infiltration into the spinal cord, consistent with reduced egress from the lymph nodes and spleen (Figure S6). Despite a 35-day prophylactic low-dose regimen, siponimod significantly altered lymphocyte and macrophage subset composition in spleen and lymph nodes (Figure S6).

**Figure 2.**
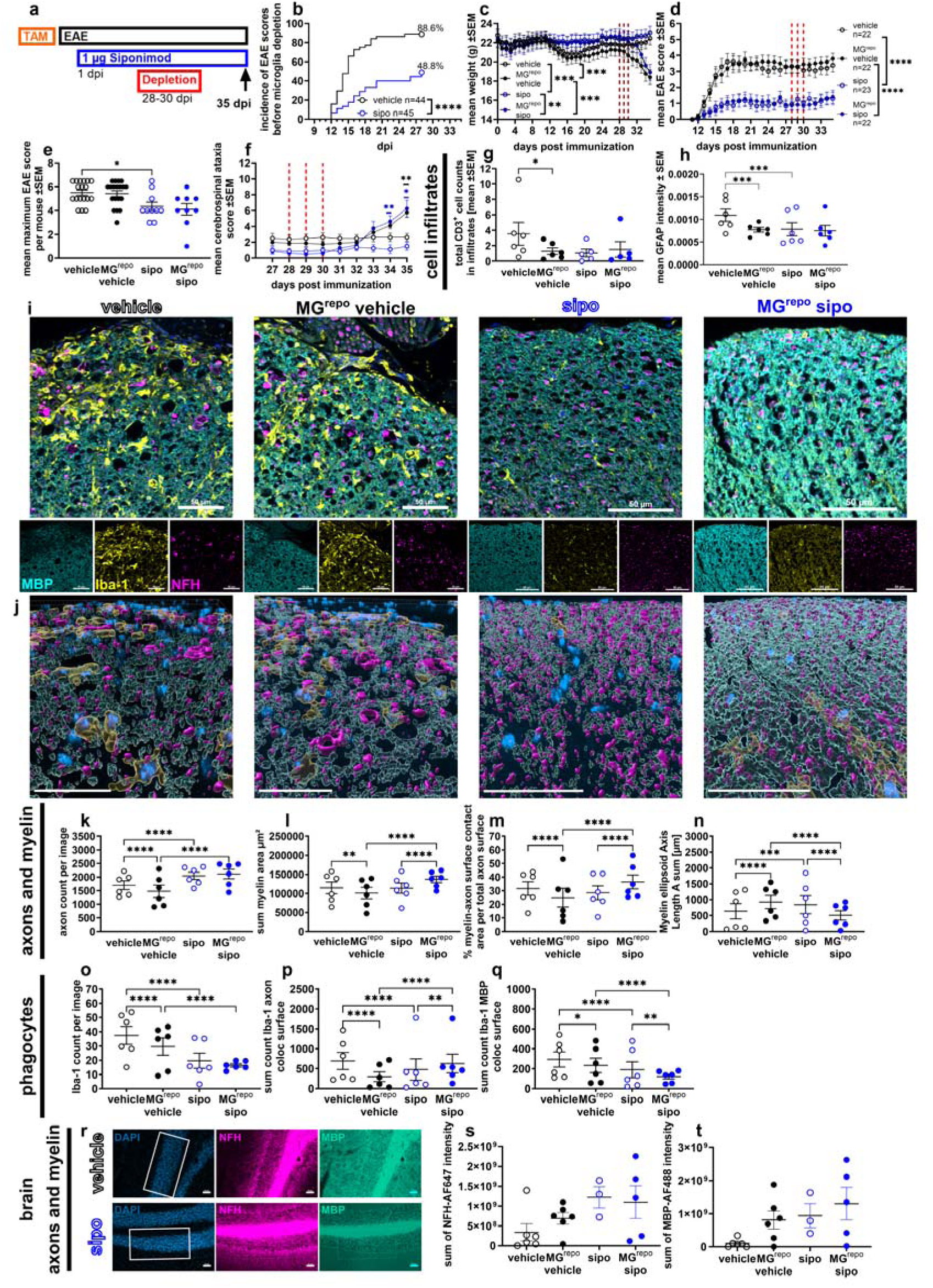
EAE and spinal cord pathophysiology alterations by siponimod and microglia repopulation. (a) experimental setting. Experimental autoimmune encephalomyelitis (EAE) with a preventive siponimod (sipo) treatment paradigm presenting incidence (b), weight loss (c) and EAE scores (d,e). Conditional microglia depletion is indicated with red lines followed by microglia repopulation (MG^repo^), leading to an atypical cerebrospinal ataxia (f) (n=8). In the (immuno-) histological stainings, the EAE pathophysiology was reflected by infiltration of CD3^+^ T cells (g), and astrogliosis (GFAP, h). Imaris-assisted analysis of spinal cord infiltrates (i,j) was used to determine axonal and myelin properties and microglia/macrophage phagocytic properties with axonal count (k), myelin area (l), myelin-axonal contact area (m), myelin ellipsoid axis (n), phagocytes (o), Iba-1 axon colocalized (coloc) surface (p), and Iba1 myelin coloc surface (q). Neurodegeneration in the brain was assessed by NFH (s) and MBP (t) staining of n=3-6 animals per group. The exemplary staining of the cerebellum (r) shows the measured area indicated with the white rectangle. Dpi – days post immunization (EAE induction), TAM – tamoxifen administration for diphtheria toxin receptor induction, NFH – neurofilament heavy (axon marker), MBP – myelin basic protein. Data are shown as mean ±SEM. Clinical data and brain immunostaining were assessed by Kruskal-Wallis test with Dunn’s multiple comparisons tests. The cerebrospinal ataxia differences per day were analyzed by 2-way ANOVA and Tukey’s multiple comparisons test. Spinal cord analyses were tested with 2-way ANOVA with Šídák’s multiple comparisons test with mean, SEM and n=5-6 per animal. Scale bars are 50 µm. The graphs are showing mean with SEM. P-values are depicted as * - p≤0.05, ** - p≤0.01, *** - p≤0.001, **** - p≤0.0001.

To better understand spinal cord pathology, we utilized three-dimensional reconstruction techniques, following staining for axons (NFH), myelin (MBP), and phagocytes (Iba-1) (Figure 2i,j). Consistent with the ameliorated EAE course, there was greater axon density with sipo treatment than with vehicle, suggestive of neuroprotection (Figure 2k). MG^repo^ reduced the number of axons compared to vehicle control, whereas sipo prevented this axonal reduction regardless of MG^repo^. As the integrity of axons is supported by intact myelin sheaths, the results for the surface contact area between axons and myelin showed the same treatment effects (Figure 2m). The demyelinated area in the white matter was significantly reduced by sipo, irrespective of MG^repo^ (Figure S5b), prompting further analysis of myelin structure. The myelin area was altered by the MG^repo^ vehicle and MG^repo^ sipo in a similar way to the myelinated axon surface (Figure 2l) with a smaller myelin diameter in the vehicle and the MG^repo^ sipo group compared to sipo and MG^repo^ groups (Figure 2n).

Iba-1^+^ cell counts were evaluated within the infiltrates, which represent both infiltrating macrophages and microglia reacting to tissue damage. The highest number of Iba-1^+^ cells was observed in areas of inflammation in vehicle-treated animals, which was significantly higher than both in MG^repo^ and sipo (Figure 2o). We found fewer Iba-1^+^ cells in MG^repo^sipo compared to MG^repo^vehicle. By calculating the colocalization surface between Iba-1+ reconstructed cells and either axon or myelin structures, it was possible to assign myelin and axonal structures within Iba-1+ cells, indicating that these cells are phagocytes. Sipo reduced the number of phagocytic structures in Iba-1 cells (Figure 2p,q). Both MG^repo^ and sipo reduced Iba-1-axon colocalization, suggesting that both reduce axon phagocytosis. In MG^repo^sipo, there were more Iba-1-axon colocalization objects compared to sipo. The Iba-1-myelin colocalization surface counts were similarly decreased after MG^repo^ and sipo. The NFH and MBP analysis of the cerebellum revealed no significant changes in the sum of the fluorescence intensity (Figure 2r-t). There was, however, an increase in fluorescence intensity in the sipo-treated groups compared to vehicle, which supports the spinal cord analyses. In summary, sipo treatment led to a reduction in spinal cord inflammation, astrogliosis, and demyelination in EAE, while preserving axonal integrity and myelination during MG^repo^. Additionally, it decreased phagocytic activity of Iba-1^+^ cells in infiltrates, thereby underscoring its neuroprotective effects.

### Sipo modulates microglia during microglia repopulation

35 days after immunization and 5 days after microglia depletion, mice were sacrificed, and brain microglia were characterized by flow cytometry to elucidate the effects on microglial states (Figure 3a). In MG^repo^ mice, microglia were starting to repopulate but had not yet fully repopulated (Figure 3b). Repopulating microglia were characterized by lower expression of homeostasis markers CX3CR1 and P2RY12 and upregulation of inflammation-associated markers CD86 and I-A/I-B (Figure 3b,c,f), supported by gMFI measurements (Fig. S7). Sipo reduced the frequency of CD86^+^ and MHC-II^+^ microglia during repopulation (Figure 3b) and, additionally, elicited upregulation of CX3CR1^+^ and CD163^+^ microglia frequencies (Figure 3c). CD206 and CD68 remained unaltered between the experimental groups.

**Figure 3.**
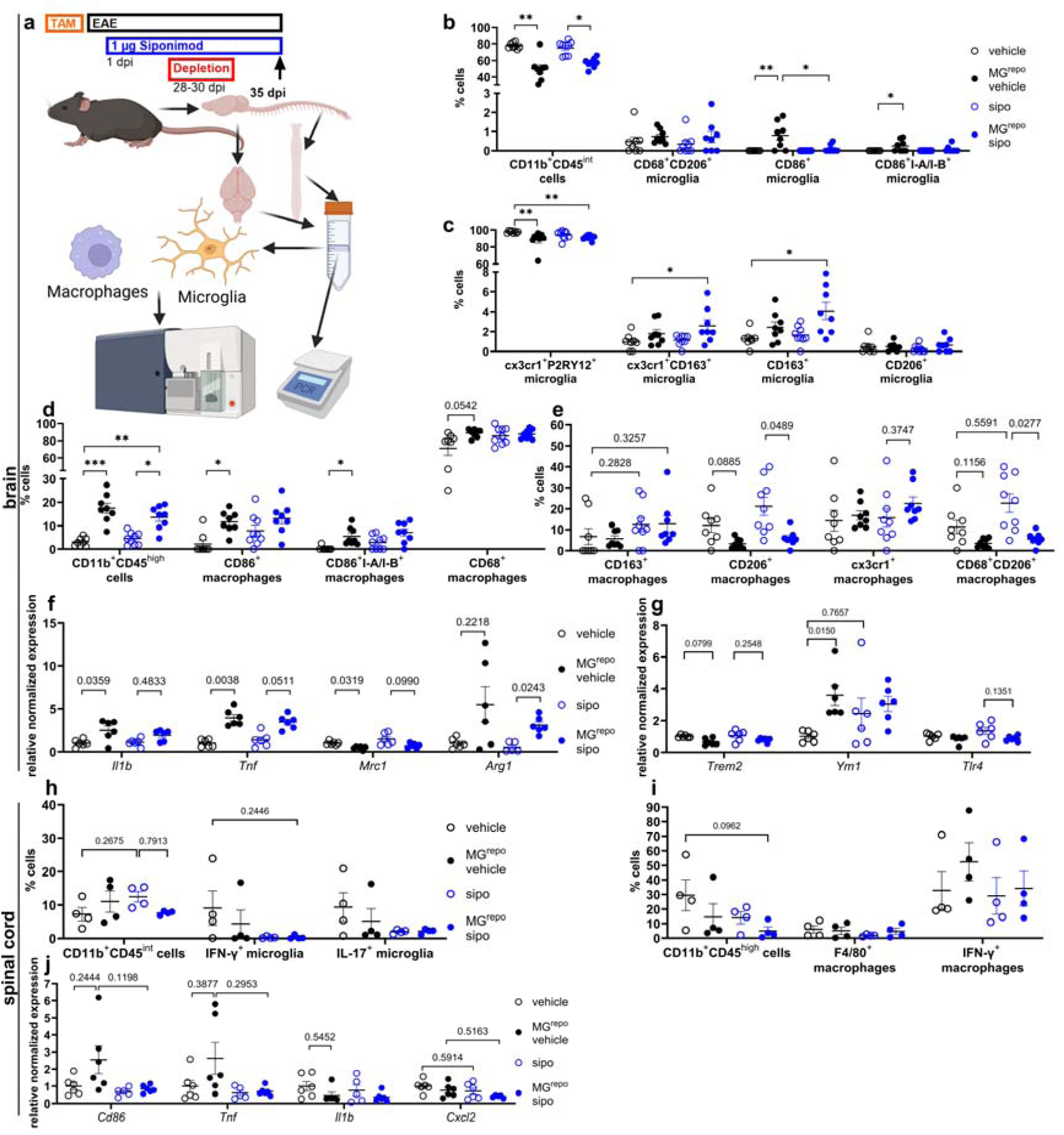
Microglia and macrophages are altered by microglia repopulation under siponimod treatment in chronic EAE. (a) experimental design of experimental autoimmune encephalomyelitis (EAE) with siponimod (sipo) treatment and timing of microglia depletion by diphtheria toxin injection and repopulation (MG^repo^) and analysis of CNS tissues. Flow cytometry (n=8-9) and qPCR (n=6) analyses of brain (b-h) of microglia (b,c) and macrophages (d,e). qPCR of brain immune cells isolated by density gradient centrifugation (f,g). The spinal cord (h-j) was analyzed by flow cytometry (h-I, n=4) and qPCR (j, n=5-6). Data are shown as mean ±SEM. Kruskal-Wallis tests with Dunn’s multiple comparisons tests. (a) was in part created with Biorender. P-values are depicted as * - p≤0.05, ** - p≤0.01, *** - p≤0.001, **** - p≤0.0001.

Furthermore, the role of infiltrating macrophages in sipo treatment was analyzed in detail. Macrophages increasingly infiltrated the brain upon the absence of microglia after depletion (Figure 3d, Figure S6j). Furthermore, the retention of macrophages in secondary lymphoid organs by sipo did not appear to be a significant phenomenon, as no substantial differences were observed between MG^repo^ vehicle and MG^repo^ sipo (Figure 3d). The infiltrating macrophages in MG^repo^ were proinflammatory despite sipo treatment with no significant changes in the expression of markers CX3CR1^+^, CD206, CD163, and CD68 (Figure 3d,e). Phagocytes expressing CD206 were lower in the MG^repo^ groups, while showing higher percentages with sipo treatment (Figure 3e). These macrophage-specific analyses provide no clear evidence for sipo-mediated modulation of macrophages in the brain.

In the brain, lymphocytes and macrophages increased after microglia depletion and presumably filled the gap left by absent microglia, as suggested by correlation analyses (Figure S6j,k,l). Of note, the infiltration of antigen-presenting lymphocytes was significantly reduced following sipo treatment (Figure S6i).

We aimed to confirm the brain immune cell alterations by gene expression analysis. We documented the same proinflammatory trend induced by MG^repo^ with elevated expression of *Il1b* and *Tnf*, and reduced expression of *Mrc1* (Figure 3g). We analyzed genes associated with the regulated phenotype and mechanisms important for brain regeneration, which could not be included in the flow cytometry analysis. *Arg1* was significantly upregulated in the MG^repo^ sipo brain compared to sipo (Figure 3g). *Trem2* expression was reduced after MG^repo^, *Ym1* expression was elevated in all groups compared to vehicle, but only the MG^repo^ group showed statistical significance, and *Tlr4* was not differentially expressed (Figure 3h).

The microglia depletion or repopulation status in the spinal cord could not be verified by flow cytometry, despite clear evidence obtained from flow cytometry analysis of the brain with the same cell isolation methods (Figure 3i). Although there were no differences in microglial and macrophage frequencies, the animals treated with sipo or MG^repo^ lacked microglia with high cytokine levels (Figure S6m,n). In conclusion, MG^repo^ during EAE induced inflammation-associated microglia with reduced homeostatic markers and elevated CD86 and MHC-II, whereas sipo treatment attenuated microglial activation, promoted regeneration-associated markers (CD163, CD206, CD68), and had limited effects on infiltrating macrophages, highlighting its potential to modulate CNS immune responses.

### MG^repo^ and sipo alter microglia-associated proteins in the spinal cord

Proteomic characterization of spinal cords from EAE mice treated with sipo or vehicle and MG^repo^ quantified over 5000 proteins, with a higher replicate variability in sipo-treated groups (Table S1). Differentially expressed proteins were identified based on relative quantification of changes in protein abundance (unpaired Welch’s t-test p-value < 0.05, see Table S1, Figure 4b). MG^repo^ resulted in the strongest effect on the proteome, reflecting the inflammatory response induced by the microglia depletion and repopulation, as evidenced by atypical ataxia scoring, cytokine elevations in the CNS by qPCR and flow cytometry. Comparing vehicle and MG^repo^ vehicle identified 472 differentially expressed proteins. Sipo treatment drastically reduced altered proteins between sipo and MG^repo^ sipo groups, indicating its ameliorative effect on EAE regardless of reactive microglia presence. Sipo had a greater effect without MG^repo^, with 283 differentially regulated proteins between vehicle and sipo, and solely 69 proteins between MG^repo^ groups, suggesting its potential to decrease high inflammatory status in EAE mice. Lastly, a quantitative comparison between vehicle and MG^repo^ identified 201 differentially expressed proteins.

**Figure 4.**
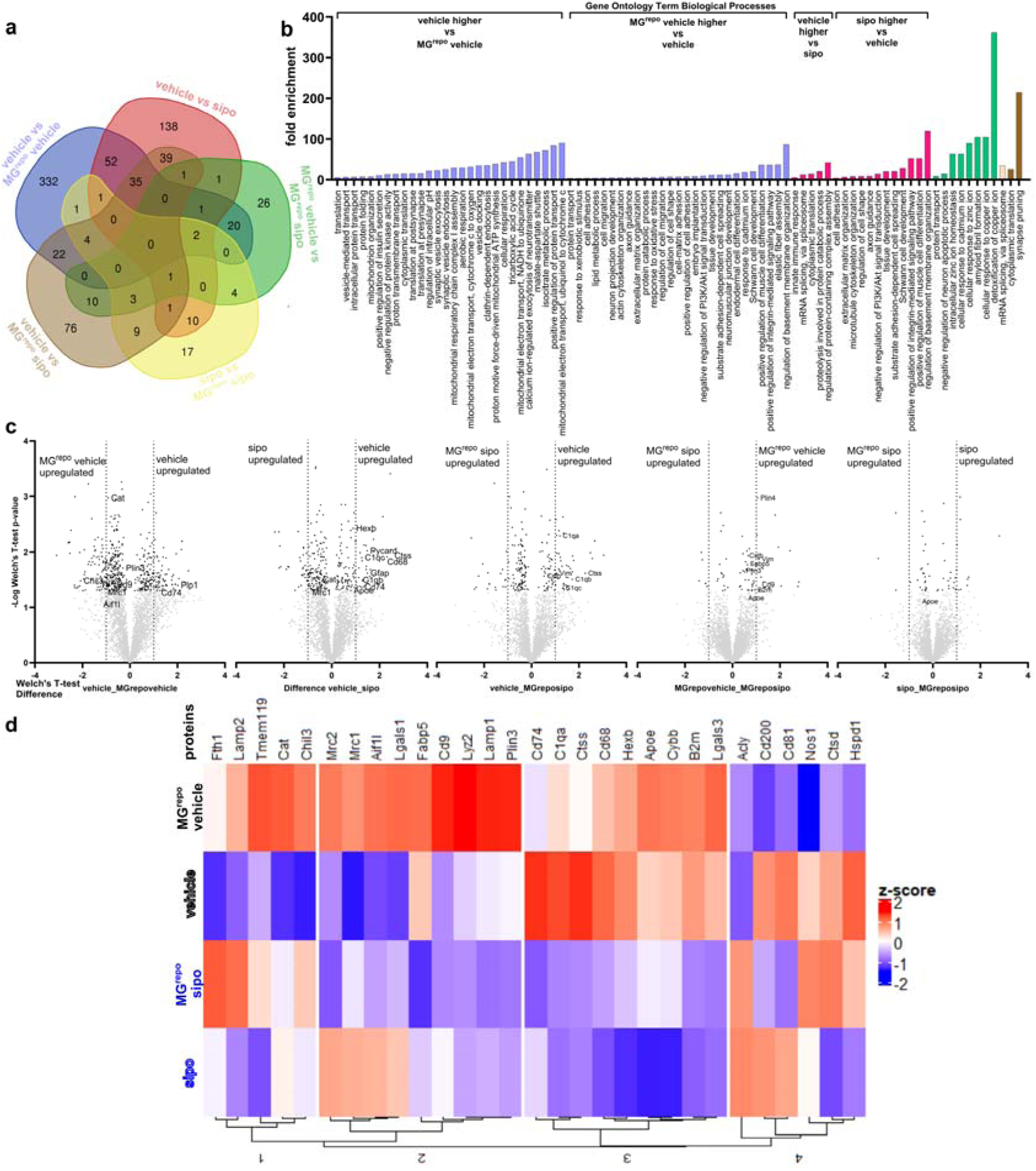
Proteomic changes in the spinal cord induced by microglia depletion and repopulation and siponimod treatment. Protein quantification by mass spectrometry (n=5) of the lumbar spinal cord tissue. The differentially quantified proteins in group comparisons are shown with matching groups between comparisons in (a). Gene ontology (GO) Term analysis on the basis of biological processes (b). The top regulated terms (p-value <0.05 and lowest Benjamini Hochberg, are displayed sorted after fold enrichment values, whereby enriched terms for proteins in vehicle mice compared to MG^repo^ vehicle mice are shown in blue with the higher abundances indicated above. Enriched terms for proteins in the comparison of vehicle mice to sipo mice are shown in red. The green bars show the enriched terms for proteins being of higher abundance in MG^repo^ vehicle mice compared to MG^repo^ sipo mice. In the yellow bar, the GO Term enriched in sipo mice compared to MG^repo^ sipo mice is presented and the brown bars show the terms enriched in vehicle mice compared to MG^repo^ sipo mice. The differentially expressed proteins in group comparisons are presented in (c) in volcano plots based on the Welch’s T test. A heatmap was created in (d) to present microglia-related protein expression differences between the experimental groups. MG^repo^ – microglia depletion with diphtheria toxin administration in cx3cr1-iDTR mice and repopulation, sipo – siponimod.

To evaluate microglial repopulation, proteomic differences between vehicle and MG^repo^ vehicle were analyzed, revealing 474 differentially expressed proteins (177 higher in vehicle and 295 higher in MG^repo^). Gene ontology (GO) term analysis based on biological processes (BP) revealed vehicle-upregulated proteins were associated with energy metabolism (TCA cycle, mitochondrial functions, synapse organization and transmission, Table S1, Figure 4b), while MG^repo^ enriched terms involved structural cytoskeleton organization, cell-matrix adhesion, and neuronal projections development.

The Top30 proteins increased in MG^repo^ vehicle vs vehicle included immunomodulatory- or oligodendrocytes- and microglia-linked proteins (Table S1). In concordance, the GO term enrichment of MG^repo^ vehicle identified a 20-fold-enrichment of the GO term “Schwann cell development” and a 6.6-fold enrichment of the term “myelination”, supporting the increased myelin diameter in histology (Table S1, Figure 2r). Additionally, several terms associated with cell structure and maintenance were enriched. Cross-comparison of vehicle vs MG^repo^ vehicle and sipo groups found only 8 differentially expressed proteins (Table S1, Figure 4a). Within the direct comparison of sipo and MG^repo^ sipo (Table S1), analyses revealed proteins with the highest abundance in MG^repo^ sipo were linked to chromatin remodeling, mRNA transcription/splicing, myelination, and immune cells, while sipo-upregulated proteins were associated with actin cytoskeleton and protein transport, with a role in oxidative stress protection.

Comparing vehicle and sipo 283 differentially expressed proteins were found (111 higher in vehicle), with GO Term enrichment analysis identifying 46 enriched terms associated with transcription and translation, antigen processing, autophagic processes, and complement activation (Figure 4c). Confirmatively, among the Top10 proteins being of highest abundance in the vehicle group, Immunity-related GTPase family M protein 3 (FC 5.6), CD68 antigen (FC 4.6), Immunity-related GTPase family M protein 1 (FC 4.4), and Interferon-inducible GTPase 1 (FC 3.4) were identified. Conversely, sipo-upregulated proteins were associated with cell structure and maintenance (membrane and ECM organization, tissue development), potentially hinting at protective processes induced by sipo treatment (Table S1, Figure 4b). Supportively, GO terms associated with neuronal cell types were enriched, among them astrocyte development, synapse assembly, synaptic vesicle transport, axon guidance, dendrite development, and Schwann cell development.

Comparison of MG^repo^ vehicle and MG^repo^ sipo yielded a small number of differential proteins (Figure 4a), indicating that microglia depletion overcomes the effects of sipo treatment. The highest fold changed proteins of MG^repo^ vehicle included Ubiquitin-like protein NEDD8 and Metallothionein-1 and 3, induced by inflammatory and stress stimuli in microglia (Table S1) (17), while MG^repo^ sipo mice exhibited increased proteasomal subunits. Lastly, vehicle vs MG^repo^ sipo comparison revealed enriched autophagy, autophagosome, and immune response-associated GO Terms in vehicle (Figure 4b), confirmed by associated high abundance proteins (Table S1). MG^repo^ sipo mice, by contrast, displayed increased abundance of proteins associated with cytoskeletal functions, protein folding, and transport. To our surprise, Periostin, a protein thought to be specifically upregulated in EAE, was of higher abundance in MG^repo^ sipo mice (18). However, Ptgds was also upregulated in MG^repo^ sipo mice. A heatmap of microglia-related proteins identified clusters with distinct expression patterns. Clusters one and two included proteins enriched in the MG^repo^ vehicle group. The clusters comprised proteins associated with homeostatic microglia (Tmem119, Mrc2), cell adhesion upregulated in MS models (CD9), phagolysosome (Lyz2, Lamp1, Lamp2, CD206), lipid droplet accumulation (Plin3, Fabp5, Cat), iron storage (Fth1), and oligodendrogenesis (Chil3) (19–27). Cluster 3 included inflammation or EAE-associated microglia proteins elevated in the vehicle groups and low differential expressions in the sipo-treated groups (CD74, Hexb, CD68, B2m, Cybb, ApoE, Ctss, C1qa, Lgals3) (21, 25, 28, 29, 20). Cluster four comprises markers for microglia (CD81), immunosuppression (CD200), antigen presentation (CD74), lipid droplet accumulation (Acly), and responsive microglia proteins associated with diseases (Ctsd, Hspd1, Nos1) (30, 31, 20, 32, 19, 21).

### RNA sequencing of the spinal cord parallels the proteomic evidence of altered microglial abundance and activation state

Given the results obtained with the global proteomics analysis, we examined the gene expression changes in the lumbar spinal cord that induced the observed alterations in myelin and microglia-associated proteins. To assess global transcriptional variance across conditions, principal components analysis (PCA) was performed. PC1 and PC2 accounted for 78% and 11% of the total variance, respectively. Samples clustered by experimental condition (naïve, EAE, and EAE MG^repo^), indicating that disease state was the primary driver of gene expression variability (Figure 5a). Consistent with the protein-level analyses, MG^repo^ displayed the most pronounced change in transcriptional profile, shifting towards the naïve state (Figure 5a).

**Figure 5.**
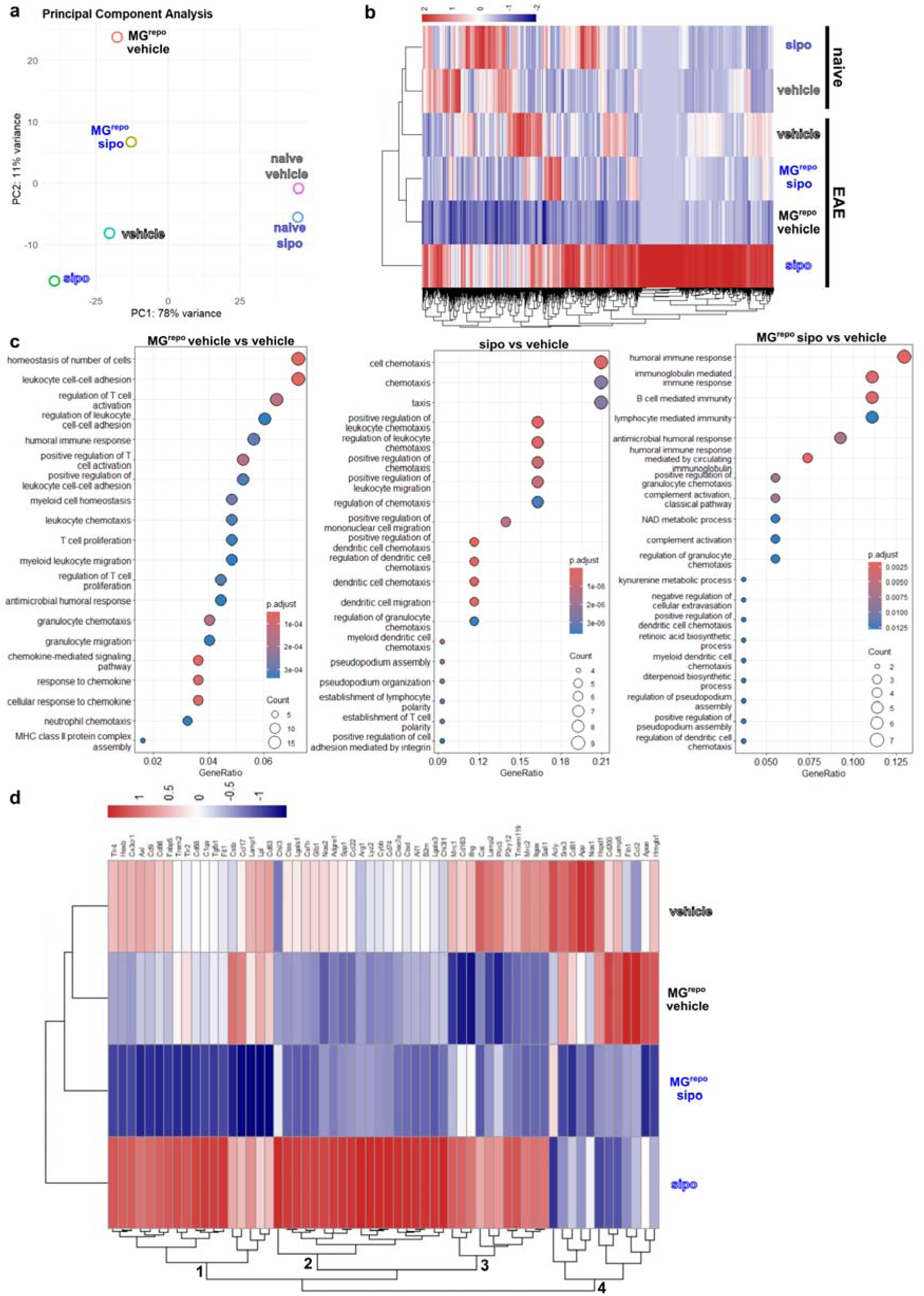
RNA sequencing of lumbar spinal cord tissue reveals effects of siponimod treatment and microglia depletion. RNA sequencing of pooled samples per group of the lumbar spinal cord tissue. DEGs in group comparisons are shown as principal component analysis (a) and heatmap (b). Gene ontology (GO) Term analysis on the basis of biological processes (c). A heatmap was created in (d) to present microglia-related DEGs between the experimental groups with EAE. PCA was conducted with normalized and transformed gene-level count data by variance-stabilizing transformation with the DESeq2 package and plotted with ggplot2. DEGs were identified in Rstudio by a p-value threshold of <0.05 with a fold change of 2 and normalized using the row-wise z-score adjustment. The heatmaps were created with the pheatmaps package. GO term analysis of BP was conducted with the clusterProfiler package with a threshold based on an adjusted p-value of <0.05 (Benjamini-Hochberg correction) and q-value <0.2, and visualized with the dotplot package. MG^repo^ – microglia depletion with diphtheria toxin administration in cx3cr1-iDTR mice and repopulation, sipo – siponimod, naïve – no EAE induction.

Global differences in gene expression were further examined using a heatmap generated using log2 fold change (FC) values. The analysis identified 3,895 genes as significantly differentially expressed (DE) with a FC (M value) of ≥ |2| (Figure 5b, Table S2). EAE sipo samples showed predominant upregulation, whereas MG^repo^, particularly in the absence of sipo, resulted in an overall downregulation of the DE genes when compared to the other experimental conditions (Figure 5b). Notably, the MG^repo^ group exhibited expression patterns closely resembling naïve samples (Figure 5b).

Differential expression analysis using NOISeq package was used due to its ability to simulate technical replicates to estimate the differential expression probability (33, 34). We identified 407 DE genes between MG^repo^ and vehicle-treated EAE spinal cords (q ≥ 0.95; |FC| ≥ 2). GO analysis revealed enrichment of immune-related BP, including B-cells and T-cell associated pathways and chemokine signaling (Figure 5c). Among significantly enriched BPs, we found upregulation of genes *Ccl8, Cxcl10, Ccr7, Ccl22, Ccr2,* and *Ccl2,* which point to enhanced immune cell migration and activation, consistent with inflammatory signaling pathways identified at the protein level (Figure 5C, Table S2). KEGG analysis revealed enrichment of the cytokine-cytokine receptor interaction (p= 0.0033) and chemokine signaling (p= 0.0055) pathways in MG^repo^ spinal cord when compared to vehicle (Figure S10).

In EAE mice treated with sipo, 161 DE genes were identified relative to vehicle controls. GO analysis revealed enrichment of pathways related to chemokine signaling and immune cell migration (Figure 5c). Significant enrichment was observed for regulation of dendritic cell chemotaxis (p= 9.96E-09), positive regulation of leukocyte (p= 1.01E-06), mononuclear cell migration (p=1.22E-06), and pseudopodium assembly (p= 1.25E-06). The Kyoto Encyclopedia of Genes and Genomes (KEGG) analysis further supported these findings, identifying significantly enriched pathways related to cytokines and chemokine signaling, including the complement and coagulation cascade (p= 0.0368), hematopoietic cell lineage (p= 0.0368), and NF-κB signaling (p= 0.0441) (Figure S10).

Microglia repopulation in the presence of sipo (MG^repo^ sipo vs vehicle) resulted in 140 DE genes (Table S2), with GO enrichment of BPs related to humoral immunity and immune cell trafficking (Figure 5c). Upregulation of genes involved in complement activation, kynurenine metabolic process (Kmo and Aldh8a1), and retinoic acid and diterpenoid biosynthetic processes (Rdh9) suggest metabolically driven immune modulation (Figure 5c), consistent with terms identified in the proteome. Comparison of MG^repo^ sipo with MG^repo^ vehicle identified 175 DE genes associated with cytoskeletal regulation and energy metabolism (Figure S10c,d), highlighting the effects of sipo in EAE animals with repopulated microglia.

To complement the microglia-specific protein expression analysis in the spinal cord, we examined a heatmap of microglia-related genes. Microglia-associated genes were upregulated in the sipo-treated EAE samples, while microglia depletion and repopulation resulted in reduced expression of canonical microglial markers, including *P2ry12*, *Tmem119* and *Sall1* (Figure 5d). This effect was more profound in the MG^repo^ sipo group and parallels the proteomic evidence of altered microglial abundance and activation state. This data supports a coordinated transcriptomic-proteomic response to microglia repopulation and sipo treatment.

## Discussion

Despite great advances over the past years to improve treatment outcomes for pwMS, the treatment of underlying progression is still unsatisfactory (1). In this study, we focused on microglial modulation in MS models to establish a neuroprotective environment following depletion of disease-associated microglia. As iron rim lesions and iron-loaded microglia are abundant in slowly expanding lesions, especially common for pMS (35), we performed iron toxicity studies (15). 1 µM sipo, resembling the amount of drug administered in humans (36, 13), was effective in both human and murine cell culture systems. The dose-dependent reduction in cell death indicates the protective properties of sipo against iron toxicity, thereby benefiting neurons and microglia. This indicates that sipo has the potential to prevent neurodegeneration in iron rim lesions in patients with multiple sclerosis (pwMS). The increased iron toxicity in the presence of microglia *in vitro* indicated a destructive microglia polarization by iron (37). Along with reduced levels of cytokine- and splenocyte-induced polarization of highly reactive microglia, we demonstrate the broad beneficial effects of sipo on CNS cells in the context of neuroinflammation, as shown before (38–40).

In EAE, we investigated treatment paradigms that should resemble the currently emphasized early treatment initiation to maximize the drug’s potential protective effect (2). The approach was to use a low dose to enable breakthrough disease, rather than prevent it completely. The objective was to potentially detect the additive effects of long-term administration and microglia depletion, and to ascertain how that affects immune cells. Despite disease amelioration with 0.45 µg sipo daily, as previously shown in C57BL/6 (36), with only a minor effect detectable on peripheral and brain immune cells. It is plausible that the initial prevention of lymphocyte egress from the lymphoid organs, presumably present at treatment initiation and reducing EAE initiation, was compensated for over the 27 days of treatment (11). Another possibility is that the prevention of lymphocyte egress was initially not significant, suggesting that sipo reduced the development of EAE by other mechanisms, such as preventing microglia and astrocytes from becoming destructive and preventing cell-cell interactions, as shown in the cell culture experiments. We intended to reset the microglia population during chronic EAE to simulate ablation of disease-associated microglia (41). The hypothesis under consideration posits that reduced peripheral immune cell numbers in the CNS during this specific phase would facilitate the repopulation of microglia that are not primed by leukocytes. Together with sipo treatment, the inflammation-associated induction of repopulating microglia should be further reduced by preventing CNS infiltration and direct modulatory effects of sipo on microglia, as shown in cell culture (3, 14). Nevertheless, the low dose of 0.45 µg sipo did not alter the microglia population.

Based on these results, we decided to implement a higher dose of 1 µg sipo to ensure sufficient CNS penetration (13), which effectively ameliorated EAE. In addition to the expected reduction in spinal cord pathology with sipo treatment, pathology was also reduced by microglial repopulation during the chronic phase of EAE (41). Microglial depletion and repopulation increased both lymphocyte and glial cell populations in the CNS; however, sipo treatment improved glial cell abundance and activation but not lymphocyte abundance or activation, in addition to mild changes in spleen and lymph node lymphocytes (13, 42). The reduced frequency of EAE-associated cells was most pronounced in sipo treatment with microglial repopulation. Microglia are abundant in EAE spinal cord lesions, so their depletion is thought to have resulted in the ablation of disease-promoting microglia within the lesions. This reduced the overall size of the infiltration by repressing cell numbers, reduced recruitment of T cells to the spinal cord due to the absence of microglia (3), and limited interaction with astrocytes and thereby lowering activation (26). Astrogliosis and microgliosis were significantly reduced by sipo treatment, supporting the hypothesis that the concentration of sipo reaching the spinal cord may be high enough to exert a direct regulatory effect on these cells (10, 38). Microglia were not completely repopulated at the time of analysis, particularly in combination with sipo treatment in the spinal cord and brain (43). The infiltration of reactive peripheral immune cells into the microglia-ablated CNS is a process in consequence of the replenishment of the redundant microglia niche, and a response to the accumulation of cell debris. The infiltrating peripheral monocytes were previously identified as inflammatory and to cause the observed atypical ataxia following microglia depletion (16). Sipo presumably reduced this infiltration, explaining the lowered microglia/macrophage-related gene regulation and protein expression detected upon MG^repo^. The microglia repopulation under sipo treatment led to altered protein expression, reducing antigen-presenting properties and inflammation-associated proteins while enhancing protein expression associated with regenerative properties (3). High biological variability within the experimental groups limited statistical analysis and data interpretation, necessitating caution when interpreting proteomic results. Myelinoprotective properties of sipo have also been described in de- and remyelination models (39, 12, 40). Sipo treatment *in vitro* and *in vivo* demonstrated neuroprotective properties against iron toxicity and prevented neuronal cell loss in the spinal cord and brain, possibly through direct effects on neurons and indirect effects by maintaining myelin integrity and modulating reactive cell types, such as microglia. In particular, microglial resetting in the chronic phase of EAE and subsequent microglial repopulation under sipo treatment were associated with beneficial effects on spinal cord pathology and microglial function.

Our study has several limitations. First, although EAE is an established and well-characterized model that enables the use of genetically modified mouse lines, it does not fully recapitulate all aspects of progressive disease (44). We implemented microglia (MG) depletion to enable robust repopulation of CNS phagocytes under divergent experimental conditions. While the CX3CR1-CreER/iDTR model is widely used and well described, our study has specific constraints. First, we did not verify microglia-restricted DTR expression by antibody-based staining. Second, in our flow-cytometry analyses we could not unequivocally distinguish repopulating microglia from surviving microglia, as we did not include a proliferation marker to identify microglia that repopulated via local proliferation. Sipo treatment was initiated at EAE induction to model early treatment onset, followed by addition of a microglia-targeting intervention after relapse. Therefore, because immune activation and neurodegenerative processes may already be influenced by sipo before microglia depletion, an additional experimental paradigm in which sipo is initiated concomitantly with microglia ablation would be important to corroborate our conclusions. To improve translational relevance, future studies should complement genetic depletion with non-genetic microglia ablation approaches, including CSF-1R antagonists (45). Moreover, single-cell approaches could help to resolve microglial and other CNS-resident or infiltrating populations at higher resolution and to validate findings derived from pooled samples.

A major strength of our work, however, is the comprehensive assessment of the model, enabling microglia-specific depletion without affecting macrophages, integrating analyses of peripheral immune cells, brain, and spinal cord. Collectively, our data support the concept that microglial modulation can exert beneficial effects in chronic neuroinflammation, with potential relevance to multiple sclerosis progression and related neurodegenerative disorders.

## Materials and Methods

### Sex as a biological variable

Our study examined female and male mice. Similar findings were reported for both sexes, unless stated otherwise. For EAE experiments, the results per experimental group were analyzed regarding sex differences using the Mann-Whitney test (Table S6).

### Cell culture of human neurons and microglia

Fetal brain-derived neurons and microglia were used from deidentified samples as previously described (46, 47). The cells were harvested after 12-15 days of culture. Neurons were harvested by trypsinization and seeded at a concentration of 10^5^ cells per well in pLorn-coated 96-well flat-bottomed black plates (Falcon 353219) following culture in medium with Ara-C over 72 hours. Thereafter, neurons were pretreated with sipo, and after one hour 2.5*10^4^ microglia were added following the addition of FeSO_4_ in saline (Figure 1a). The co-culture was conducted for 24 hours with live cell imaging by acquiring propidium iodide (PI, ThermoFisher Scientific) and Hoechst 33342 NucBlu Live ReadyProbes reagent (ThermoFisher Scientific) signal every two hours with a Celldiscoverer 7 (Zeiss) or an ImageXpress Micro XLS High-Content Analysis system (Molecular Devices). The PI and Hoechst33342 positive cells were analyzed using ZEN software (Zeiss) or MetaXpress (Molecular Devices).

### Cell culture of murine neurons and microglia

The C57BL/6J pregnant mice, postnatal mice, and embryos were sacrificed by methods that comply with European and German animal welfare guidelines prior to tissue explants. The cerebral hemispheres of embryonic day 15-16 mouse embryos were dissected to obtain neurons as described before (46). 10^5^ neurons were seeded in 100 µL B-27 Plus Neuronal Culture System medium (Gibco A3653401) in a pLorn-coated 96-well plate and cultured for 48 hours, adding 100 µL Neurobasal medium (Gibco 21103049) after 24 hours.

Primary murine microglia were prepared from postnatal day 0-2 C57BL/6 mice. Cortices were mechanically dissociated and seeded into Poly-D-lysin hydrobromide-coated (10 µg/mL PDL; Sigma-Aldrich P6407-5MG) cell culture flasks in supplemented DMEM medium (Table S6). Microglia were harvested as they floated over the astrocyte layer. After seeding, the medium was exchanged after 1 hour to Neurobasal medium.

To isolate splenocytes, spleens of adult mice were mechanically dissociated and separated via density gradient centrifugation with Ficoll® Paque Plus (Cytiva 17-1440-02) as described before (36). The cell pellet was resuspended in 1 mL of complete neurobasal medium (Table S6). For stimulation, splenocytes were incubated in medium supplemented with PMA (1 μg/mL) and ionomycin (iono, 20 ng/mL) for 4 hours. For co-culture with 3*10^4^ primary microglia, 5*10^4^ splenocytes were added into each well (Figure S2k) in complete neurobasal medium.

For assays of neuron-microglia co-cultures, neurons were pretreated for one hour with sipo (Hycultec HY-12355 10 mM in DMSO) in concentrations measured previously in the CNS after oral administration (12) or DMSO as a control. Meanwhile, microglia were harvested. 2.5*10^4^ microglia were added per well onto pretreated neurons. After one hour, a full medium change was conducted (Figure 1h). The same concentration of sipo was added again with the stimulating agents like FeSO_4_ (Carl Roth P015.1 solubilized in 0.9% NaCl), or IFN-γ (BioLegend 575302, 30 ng/mL;i) and LPS (Sigma-Aldrich L4391, 1 µg/mL) (iLPS). To investigate the inhibition of stimulation by sipo, microglia were pretreated with sipo for one hour before adding FeSO_4_, IFN-γ, LPS, or splenocytes.

The toxicity of sipo, FeSO_4,_ and iLPS was assessed by live cell imaging with Hoechst 33342 (Thermo Fisher Scientific H3570) and PI (Invitrogen 00-6990-50) before imaging every two hours in the Cytation5 (Agilent BioTek) over the course of 24 hours. The analysis of positive cells was done automatically with the Gen5 software (BioTek Instruments).

For immunocytochemistry (ICC), PFA-fixed 96-well plates were permeabilized with 0.2% Triton X-100 for 10 minutes and blocked with 5% goat and donkey serum each in 0.2% Triton X-100 for 1 hour. Primary and secondary antibodies (see Table S3) were incubated in blocking solution. Eight images per well were acquired using the AxioObserver microscope (Zeiss) and analyzed with the ZEN blue 3.10 software (Zeiss). For flow cytometry, the cells were detached using Accutase (Invitrogen 00-4555-56) and transferred into a 96 well V-bottom plate for staining and measurement. Cell surface markers were labelled with antibodies (see Table S3) for 30 minutes at 4 °C, making use of Fc-blocking and True-Stain Monocyte Blocker™ (Biolegend) to increase specificity. Viability was determined using Zombie Aqua™ Fixable Viability Kit (Biolegend) according to the manufacturer’s instructions. Detection of fluorescence signals was achieved with BD FACSCelesta™ Flow Cytometer and analyzed with FlowJo (FlowJo X 10.0.7r2, BD). These methods further enabled routine assessment of culture purity via immunolabeling with microglial markers.

### Experimental autoimmune encephalomyelitis

All animal experiments were approved by the animal care committee of North Rhine-Westphalia, Germany (AZ 81-02.04.2019.A425). Mice of both sexes aged 8-12 weeks were included. To induce specific microglia ablation during EAE, we crossed homozygous B6.129P2(C)-*Cx3cr1^tm2.1(cre/ERT2)Jung^*/J (The Jackson Laboratory 020940, cx3cr1) and homozygous C57BL/6-Gt(ROSA)26Sor^tm1(HBEGF)Awai^/J (The Jackson Laboratory 007900, iDTR) to generate DTR^MG^ mice. Mice were housed under environmentally controlled conditions and a 12:12 h dark-light cycle under pathogen-free conditions at the central breeding facility of the Medical Faculty of Ruhr-University Bochum. Mice had access to chow and water *ad libitum*. EAE was induced as described before (36). Animals were scored and weighed daily according to a previously defined scoring scheme (36). Mice with a score of 7 were euthanized according to animal care guidelines. For each mouse, all scores per day in each experimental group were included in the statistical testing with the Kruskal-Wallis test followed by Dunn’s post hoc multiple comparisons. Before treatment initiation, animals were randomized according to weight or score in the therapeutic experiments.

On the day after EAE induction, the animals were divided into 2 groups and given either rapeseed oil (vehicle) or sipo (Hycultec HY-12355) dissolved in rapeseed oil by oral gavage. To specifically ablate microglia, first the DTR^MG^ mice were administered 1 mg of tamoxifen (Sigma-Aldrich T5648-1G; TAM) in corn oil i.p. every other day for five days at weaning age to induce diphtheria toxin receptor (DTR) expression on cx3cr1-expressing cells. 1 µg diphtheria toxin (Sigma-Aldrich D0564-1MG; DT) injection i.p. for three days (MG^repo^) was conducted at least four weeks after TAM injection. After microglia ablation by DT administration, the cells begin to repopulate approximately five days after the last DT injection (43). The mice were sacrificed as described. Whole blood, spleen, lymph nodes, spinal cord, and brain were explanted and either directly processed for flow cytometry or stored in PBS or PFA.

### Flow cytometry

Cells of blood, spleen, and lymph nodes were isolated as previously described (36). Single cell suspensions were transferred to 96-well V-bottom plates, and staining (Table S3) and flow cytometry were conducted as described with the FoxP3/Transcription factor set (Invitrogen ref.00-5521-00) according to manufacturer’s instructions (36). The brain and spinal cord were dissociated mechanically. The single cell suspension was separated using Percoll gradient (70%, 37%, 30%, PBS). Cells in the interphase of 37% and 70% Percoll were stained for flow cytometry (Table S4) or pelleted and frozen for mRNA extraction and subsequent qPCR. Flow cytometry was performed using the BD Celesta (see cell culture) and the Cytoflex LX (Beckman Coulter). Gating (Figures S7-S9) and analysis were done with FlowJo software (BD). tSNE plots were created as described before (48).

### Histological analyses

All sections of different organs or section methods were selected at random based on section quality, without knowledge of experimental group, EAE scores or other analytical parameters. For the following staining protocols, the 5 µm paraffin-embedded sections were melted at 60 °C for two hours before deparaffinizing in Roti-Histol and rehydration. The spinal cord of EAE mice was stained with H&E and luxol fast blue (LFB) double staining (27 replicates per mouse: three paraffin sections with three lumbar, thoracic, and cervical parts each). Pictures were acquired with a Axio.Observer.Z1/7 microscope (Zeiss), using the tile function with a 20x objective. Measuring of white matter and demyelinated or infiltrated areas was conducted in ImageJ.

For immunohistochemical staining of the spinal cord of EAE mice, antigen retrieval was achieved by boiling in citric buffer and incubation with ice cold methanol. Primary antibodies were incubated in blocking solution overnight at 4°C (Table S3). The secondary antibodies were incubated for 1-1.5 hours in blocking solution. If necessary, the ready ProbeTM Tissue Autofluorescence Quenching Kit (Thermo Fisher Scientific) was applied for ten minutes before embedding with DAPI Fluoromount (Biozol). Pictures were acquired using the tile function and detailed pictures of infiltrates were acquired using z-stacks. Higher resolution images were acquired with apotome. Measuring the intensity of AF647 (GFAP) over the whole cross section of the spinal cord and counting the DAPI^+^CD3^+^ cells in the infiltrates was performed with ZEN software (3.10 Zeiss; 18 replicates per mouse: two paraffin sections with three lumbar, thoracic, and cervical parts each). Three-dimensional reconstruction of Iba-1, NFH, and MBP signals was conducted of local bleaching corrected apotome z-stacks with IMARIS software (10.2.0 Bitplane AG). Construction of surfaces was done by background subtraction and size limitation of the surfaces. The interactions of the surfaces were calculated with the MATLAB extensions with surface functions for surface-surface colocalization (coloc) between the NFH and MBP, Iba-1 and NFH, and Iba-1 and MBP surfaces, and for the surface-surface contact area between NFH as primary surface and NFH-MBP coloc surface as secondary coverage surface. The white matter infiltrates of thoracic and lumbar/sacral sections of two slices were analyzed (varying number of replicates: one each with 14 and 35, 11 with 28, and 10 with 24 replicates).

The PFA-fixated brains were dehydrated in 30% sucrose and embedded for cryopreservation. 20 µm sections of the brain were subjected to ascending and descending ethanol series and washed with 0.3% Triton X-100 in PBS and incubated for blocking and antibody incubation in 0.5% Triton X-100 in PBS. The anti-NFH and anti-MBP antibodies were incubated overnight, and the secondary antibodies for two hours before embedding in DAPI Fluoromount (Table S3). Images of the cerebellum were acquired with a 20x objective in z-stacks. Using the ZEN software, the z-stacks were aligned and an orthogonal projection applied. With the analysis function, a same-sized frame was analyzed of the cerebellum for NFH and MBP fluorescence intensity. The mean of replicate measurements per animal was calculated and used for statistical analysis and graphing.

### RNA isolation

The immune brain cells for qPCR were isolated by Percoll-density gradient in the same way as for flow cytometry. Brain immune cells in the interphase were pelleted, liquid removed and stored at -80 °C until mRNA isolation using the RNeasy plus mini kit (Qiagen 74136) according to manufactureŕs instructions.

Tissue pieces for RNA isolation were frozen in liquid nitrogen and stored at -80 °C until isolation. For the spinal cord, a one-centimeter piece of lumbar spinal cord was used. The tissue was disrupted with trizol and the TissueLyser II system (Qiagen) followed by phase separation with chloroform and Phase lock gel tubes (VWR 733-2478). The remaining mRNA isolation was done with the RNeasy plus mini kit.

For RNA sequencing, the RNA samples (three of each group) of lumbar spinal cord tissues were pooled to reach the needed concentration. This was necessary due to insufficient RNA yield after isolation; individual samples did not provide enough material for quality control and library preparation. As a result, three samples per group were pooled. Regardless, the samples provided good quality, and quantity reads to continue with downstream analysis of the data.

### Reverse transcriptase qPCR

cDNA transcription, qPCR and analysis were done as described before (36). The primers used are listed in Table S4. The CT values were normalized to the housekeeping genes, and the fold change expression was normalized to the control group (vehicle for brain immune cells and spinal cord).

### RNA sequencing of lumbar spinal cord tissue

Gene expression profiling of the pooled samples described in the RNA isolation section using RNA-Seq was performed by the University of Iowa Genomics Division using manufacturer-recommended protocols. The detailed methods for the global analysis of the pooled samples are provided in the supplement.

### Protein isolation and proteome analysis

In brief, the protein analysis of the lumbar spinal cord tissue samples was conducted as follows: the proteins were isolated using a urea buffer and sonication. Then, the proteins were digested to remove lipids, dried, and resuspended in TFA to enable mass spectrometry measurements. The methodological details can be found in the supplement.

### Statistics

The cell culture experiments were measured in at least triplicates per experimental group and a single experiment. All statistical analyses were performed with Prism 10 (GraphPad Software). In the experiments with replicates per experiment or animal, the mean and standard error of the mean (SEM) were calculated and used together with the N (number of replicates; see the respective methods sections) for each experiment or animal used for grouped analysis 2-way ANOVA with Šídák’s multiple comparison test. Data with one measurement (flow cytometry, qPCR, EAE score, body weight and brain histological staining) were assumed to be non-Gaussian and analyzed using the Kruskal-Wallis test with Dunn’s post hoc multiple comparisons. The EAE incidences were analyzed with survival curve comparison Log-rank (Mantel-Cox) test. Correlations were analyzed with Spearman correlation. The graphs depict the means ± SEM. Statistical significance was reached with p≤0.05 and displayed as follows: *p<0.05, **p<0.01, ***p<0.001, ****p<0.0001.

### Study approval

All animal experiments were approved by the animal care committee in Düsseldorf North Rhine-Westphalia, Germany (LANUV, no. 82-02.04.2019-A425). Primary human cells were isolated from deidentified samples in accordance with the ethics approval of the University of Calgary ethics committee(47).

### Data availability

Values for all data points in graphs are reported in the Supporting Data Values file. All data are available from the corresponding author upon reasonable request.

## Author Contributions

NH designed, performed and analyzed experiments, designed figures and wrote the first draft of the manuscript. SaF, LR, BE, A-CG, HHH, S-MO, KK, RH, DM, AK, HM, VB, CPS, GT, MVM performed experiments, acquired and analyzed data and critically revised the manuscript. SR, ME, JRP, OG, KFW, KMA, IS, VWY, RG analyzed data and critically revised the manuscript. SF designed and supervised the study, analyzed data, drafted figures and wrote the manuscript. RG and SF acquired funding. All authors read and approved the final version of the manuscript.

## Acknowledgments

The authors thank Xiomara Pedreiturria for technical support and organization in the laboratory in Bochum and Claudia Silva, Cenxiao Li, and Marlene Thorsen Mørch for technical support in Calgary. We thank the Hotchkiss Brain Institute Advanced Microscopy Platform Facility at the University of Calgary for their help. RNA-Seq data presented herein were obtained at the Genomics Division of the Iowa Institute of Human Genetics (RRID: SCR_023422), which is supported, in part, by the University of Iowa Carver College of Medicine. KFW is supported by the German Research Foundation (RTG 2862, project number 492434978 and Germany’s Excellence Strategy - EXC 2033 - 390677874 – RESOLV). We thank the Core Unit for Bioinformatics (CUBiMed.RUB) at the Medical Faculty of the Ruhr-University Bochum, Germany, for statistical support in proteome data analysis.

