## Supplements file for "Modulation of repopulating microglia in multiple sclerosis models with implication for neuroprotection"

**This PDF file includes:**

Supporting text for spinal cord methods.

Figures S1 to S10

Legends for Tables S1, S2 and S6

Tables S3 to S5

SI References 1 to 15

### Supporting Information Text

#### Detailed methods for Multiomics analysis of the lumbar spinal cord.

RNA sequencing of lumbar spinal cord tissue. Briefly, 125 ng of total RNA was used to prepare sequencing libraries using the Illumina stranded total RNA prep, ligation with ribo-zero plus library kit (Cat. #20040529, Illumina, Inc., San Diego, CA). The molar concentrations of the resulting indexed libraries were measured using the 2100 Agilent Bioanalyzer (Agilent Technologies, Santa Clara, CA) and combined equally into a pool for sequencing. The concentration of the library pool was measured using the Illumina Library Quantification Kit (KAPA Biosystems, Wilmington, MA) and sequenced on the Illumina NovaSeq 6000 genome sequencer using 100 bp paired-end SBS chemistry.

The data generated were analyzed using the Nextflow pipeline nf-core/rnaseq v3.19.0 (1), run on the Argon High-Performance Computing (HPC) cluster at the University of Iowa. Briefly, fastq data were merged and strandedness was automatically detected using Salmon Quant. The tool Trim Galore! was used to perform quality and adapter trimming of the fastQ files, then STAR was used to map the raw reads to the reference genome GRCm39 and Salmon to quantify the data. The remainder of the bioinformatic analysis was performed using Rstudio 2025.05.1 (Posit Software PBC, Boston,MA) (2).

The RNA-seq gene-count level data were imported into Rstudio where differentially expressed genes (DEGs) were identified using the statistical thresholds of p-value <.05 and fold change of absolute 2. DEGs gene expression values were extracted and normalized using row-wise z-score adjustment to facilitate the comparison among genes. The heatmap was generated using the pheatmap package (3), using hierarchical clustering to both samples and genes. Additionally, a curated list of microglia-related genes was used to filter the data set. Gene expression values for four of the experimental conditions (EAE_vehicle, EAE_sipo, EAE_MGrepo vehicle, EAE_MGrepo sipo) were extracted, normalized and visualized as described above.

For Principal Component Analysis (PCA) gene-level count data were normalized and transformed using a variance-stabilizing transformation. The DESeq2 package was used to conduct the PCA (4). The first two principal components were visualized to assess sample clustering and variance. The plot was generated using ggplot2 (5). The NOISeq R package was used as a non-parametric approach for differential expression analysis (6, 7). The NOISeq method was chosen since it was developed to deal with datasets with no replicates. The data was normalized to TMM. The data was filtered with the thresholds of log2FC ≥2 and a probability of ≥0.95, considering any gene that passes such a threshold as differentially expressed and significant. Ensemble gene identifiers were converted to EntreZ IDs and MGI symbols with the biomaRT package (8). Gene ontology (GO) and pathway enrichment analyses were performed and visualized using the clusterProfiler package (9, 10). The enrichGO() function was utilized to identify GO terms that are significantly enriched. The genes were annotated with the Bioconductor – org.Mm.eg.db (11), the mouse genome annotation database, focusing on the Biological Process (BP) ontology. An adjusted p-value of < 0.05 (Benjamini-Hochberg correction) and q-value < 0.2 were used as a significance threshold. The top 20 enriched GO terms were visualized using the dotplot package (12).

Protein isolation and proteome analysis. 50µL of urea buffer (30 mM TrisBase, 2 M Thiourea, 7 M Urea) was added to isolated spinal cord samples. Lysis was achieved by homogenization utilizing a pistil and a subsequent sonication cycle, consisting of 30 seconds sonication and 30 seconds rest on ice, in a sonication bath. Protein concentration was determined by Bradford assay and protein digestion was performed utilizing the single-pot solid-phase-enhanced sample preparation (SP3) method (13) to remove interfering lipids. Resulting peptides were dried in vacuum and resuspended in 0.1% TFA.

200 ng of each sample was injected and concentrated on a precolumn (PepMap Neo C18 Trap Cartridge 300 μm × 0.5 cm, particle size 5 μm) using the Vanquish Neo UHPLC system (Thermofisher Scientific, Bremen, Germany) and then separated on an analytical column (DNV PepMapTM Neo, 75 μm x 150 mm, C18, particle size 2 μm, pore size 100 Å). The peptides were separated at a flow rate of 400 nl/min and a solvent gradient of 1 % B to 21 % B (B: 84 % acetonitrile, 0.1 % formic acid) for 70 minutes followed by an increase in the gradient up to 40%B for 15 min. The column was rinsed with 95 % B for 5 min. The chromatography was online coupled to an Orbitrap Fusion Lumos mass spectrometer (ThermoFisher Scientific, Bremen Germany). Before entering the mass spectrometer, the peptides were ionized by ESI. The capillary temperature was set at 275 °C and the spray voltage at 1800 V. MS1 spectra were recorded in the range of 350 to 1400 m/z with a resolution of 120,000 at 200 m/z (normalized AGC target 100%, maximum injection time 54 ms). For data-independent acquisition, 20 equally sized isolation windows between 400 and 900 m/z were scanned at the MS2 level. The resolution of the Orbitrap was set to 30,000 and fragment ions were generated by HCD at a normalized collision energy of 32. The AGC target was set to 1000% and injection time to 54 ms.

The generated DIA data (raw format) were analyzed in the SpectronautPulsar program (version 19.5.241126.62635, Biognosys, Schlieren, Switzerland) utilizing the direct DIA option. The analysis settings were selected according to the manufacturer's instructions, with the exception that trypsin was added as digestion enzyme and carbamidomethylation on cysteine was set as fixed and oxidation on methionine as variable modification. The mouse reference proteome from Uniprot (accessed 03.2025, 54,739 entries) was selected as database. The abundances (MaxLFQs) of protein groups were LOESS-normalized and analyzed using R (version 4.5.0, R Core Team, 2021, https://www.R-project.org/) with the 'ProtStatsWF' package (version 1.0.0, [https://github.com/mpc-bioinformatics/ProtStatsWF]). Label free quantified (LFQ) values were further utilized for statistical analysis in Perseus (14) . Stringent filtering criteria were applied, whereby only proteins present in at least 3 replicates per group were chosen for further statistical evaluation. Remaining missing values were imputed from normal distribution utilizing a width of 0.3 and a down shift of 1.8. Welch’s t test was conducted to identify significantly differentially expressed proteins. A protein with a p-value <0.05 was considered statistically significant. To calculate the direction of regulation of a protein between two sample groups averaged log2 LFQ values per protein were subtracted from each other, resulting in a fold change (FC). Gene ontology enrichment analysis was performed as described before (15). In detail, proteins found to be of higher abundance within one comparison were functionally annotated towards biological processes (BP), as background the whole mouse proteome was chosen. A Benjamini-Hochberg adjusted p value < 0.05 was used as a significance threshold and fold enrichment scores were taken to assess the enrichment of resulting GO-terms. Unsupervised hierarchical clustering analysis was applied to proteins related to microglia with the aim of grouping the proteins with similar abundance profiles across different conditions. To this end, the mean MaxLFQs for the replicates within conditions were calculated and utilized to calculate the z-scores for each protein. The results were then visualized using a heat map. The distance metric was based on the Pearson correlation coefficient.

Figures

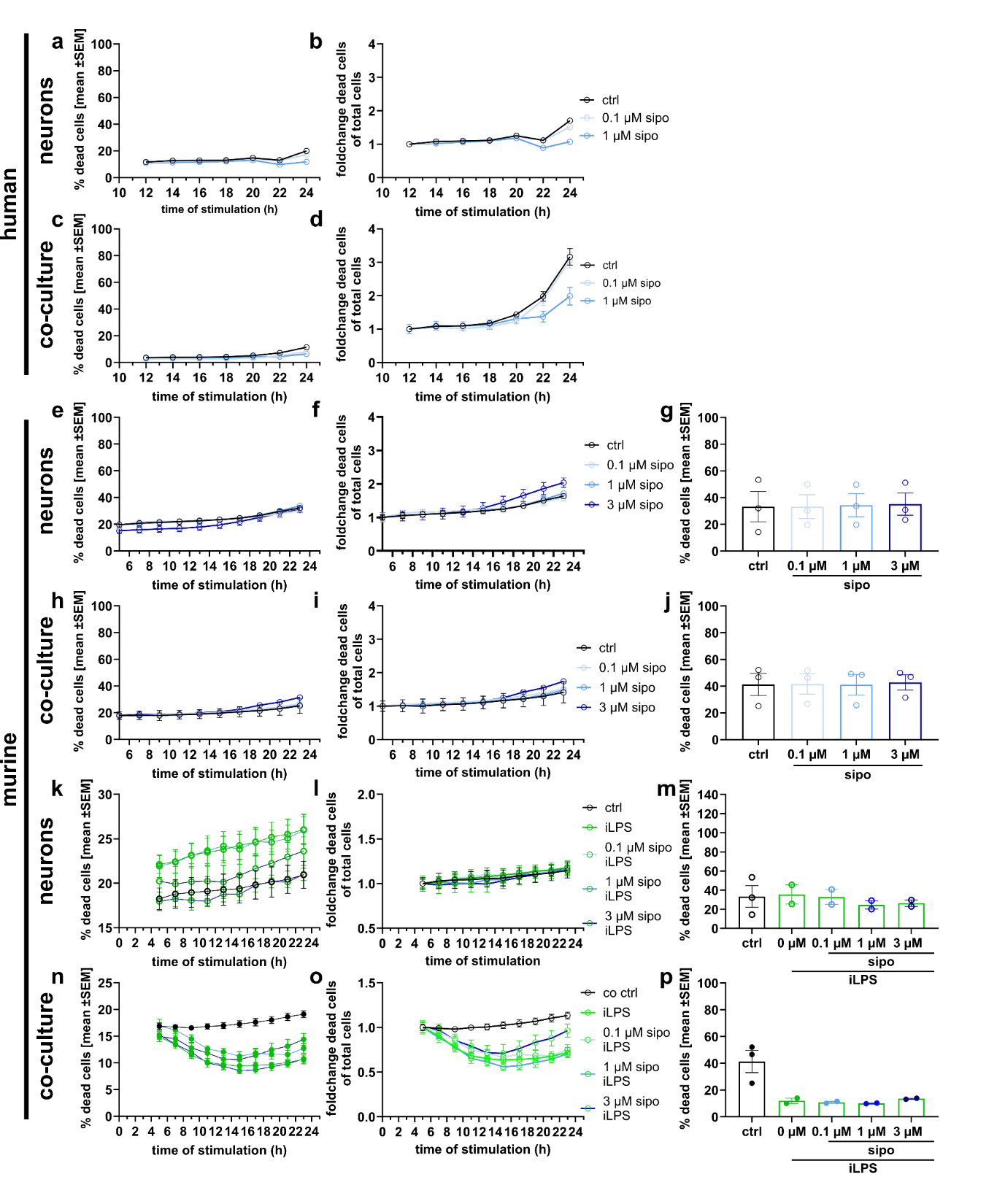

**Fig. S1.** **Siponimod and Interferon-gamma/lipopolysaccharide treatment have no adverse effects on primary human and murine neuron and microglia cultures.** Live cell imaging of control groups in human (a-d) and murine (e-p) neuron and microglia cultures treated with siponimod (a-j) or iLPS (k-p). The group comparison was performed after 22 hours after treatment start of two to three independent experiments (g, j, m, p). Co-culture – neurons and microglia cultured together, sipo – siponimod, iLPS – interferon-gamma and lipopolysaccharide stimulation, co-culture – neuron and microglia culture, h – hours, ctrl – control. Experiments were measured in triplicates and analyzed with mean, SEM and n per experiment with the 2-way ANOVA and Šídák's multiple comparisons test.

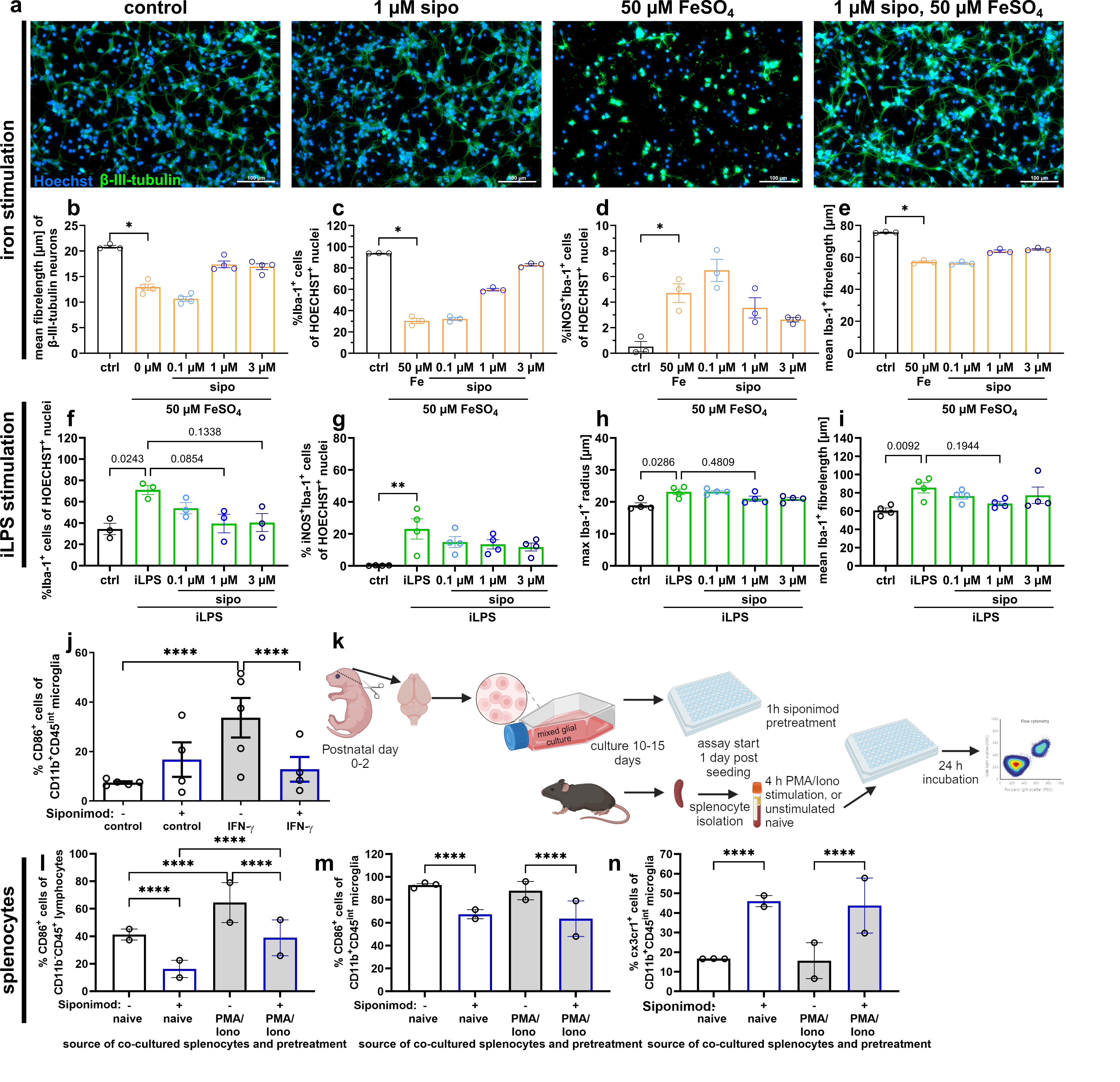

**Fig. S2. Siponimod is neuroprotective and reduces stimulation of microglia**. Immunocytochemistry and flow cytometry assessment of murine primary neuron and microglia cells. Neurite length analysis (a,b) and microglia modulation analysis in reactive marker expression and morphology (c-i) shown as one experiment measured in triplicates. Flow cytometry analysis of microglia and splenocytes (j-n) to assess microglia stimulation status by interferon-gamma and splenocytes and of splenocyte stimulation by phorbol 12-myristate 13-acetate (PMA) and ionomycin (Iono). 4-5 experiments (j) and 2-3 experiments (l-n) are presented measured at least in triplicates and analyzed with mean, SEM and n per experiment with the 2-way ANOVA and Šídák's multiple comparisons test. Data are shown as mean ±SEM. (k) was created with Biorender. P-values are depicted as * - p≤0.05, ** - p≤0.01, *** - p≤0.001, **** - p≤0.0001. Scale bars in (a) are 100 µm.

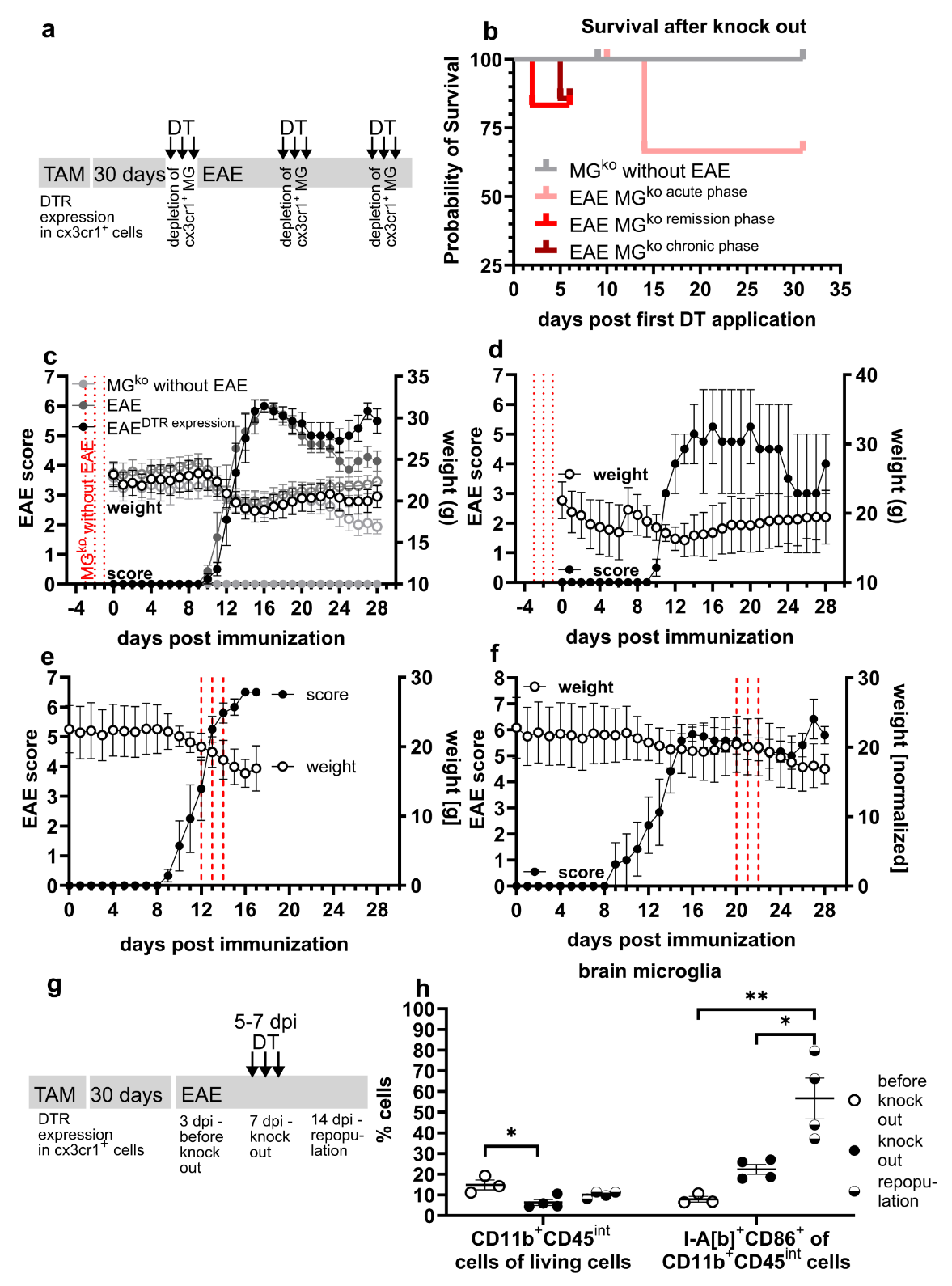

**Fig. S3. control experiments for MG^ko^-EAE and microglia depletion test.** (a-f) experiments to test adverse effects of microglia depletion (MG^ko^, indicated with red dotted lines as days of diphtheria toxin (DT) injections) at different time points in relation to experimental autoimmune encephalomyelitis (EAE) induction determining best tolerability after the acute phase of EAE. (b) n=6-7, (c) n=6-7, (d-e) n=7, (f) n=6. (g, h) microglia depletion efficacy via flow cytometry and analysis of repopulating marker expression of n=3-4 mice. DTR – diphteria toxin receptor, TAM – tamoxifen administration for diphtheria toxin receptor induction, MG – microglia, dpi – days post immunization. Data are shown as mean ±SEM. Statistical significance was tested with Kruskal-Wallis test with Dunn’s multiple comparisons. P-values are depicted as * - p≤0.05, ** - p≤0.01, *** - p≤0.001, **** - p≤0.0001.

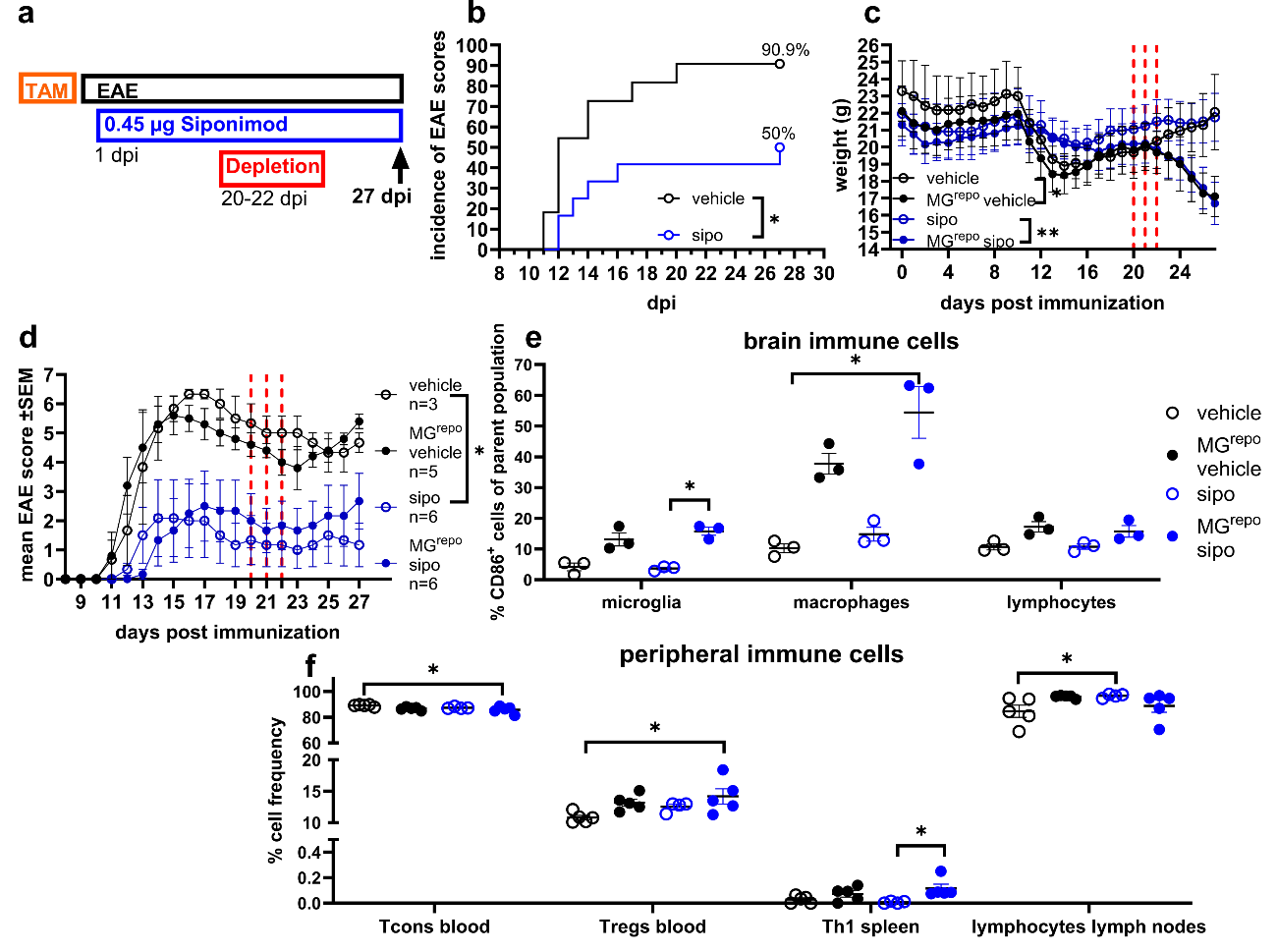

**Figure S4. Effect of low dosage preventive siponimod administration in MG^repo^-EAE.** Experimental autoimmune encephalomyelitis (EAE) experimental setup (a), incidence (b), weight (c) and clinical score (d) with flow cytometry immune cell analysis of brain (e), blood, spleen and lymph nodes (f) in mice with conditional microglia depletion on three days indicated in red. Dpi – days post immunization (EAE induction), MG^repo^ – microglia depletion with diphtheria toxin administration in cx3cr1-iDTR mice, sipo – siponimod, TAM – tamoxifen administration for diphtheria toxin receptor induction (iDTR), Tcons – conventional T cells (living CD3^+^CD4^+^FoxP3^-^ cells), Tregs – T regulatory cells (living CD3^+^CD4^+^FoxP3^+^ cells), Th1 – T helper 1 cells (living CD3^+^CD4^+^FoxP3^-^Interferon-gamma^+^ cells). Kruskal-Wallis with Dunn’s multiple comparisons tests. (b) n=11-12, (c) n as shown in (d), (e) n=3, (f) n=4-5. P-values are depicted as * - p≤0.05, ** - p≤0.01, *** - p≤0.001, **** - p≤0.0001.

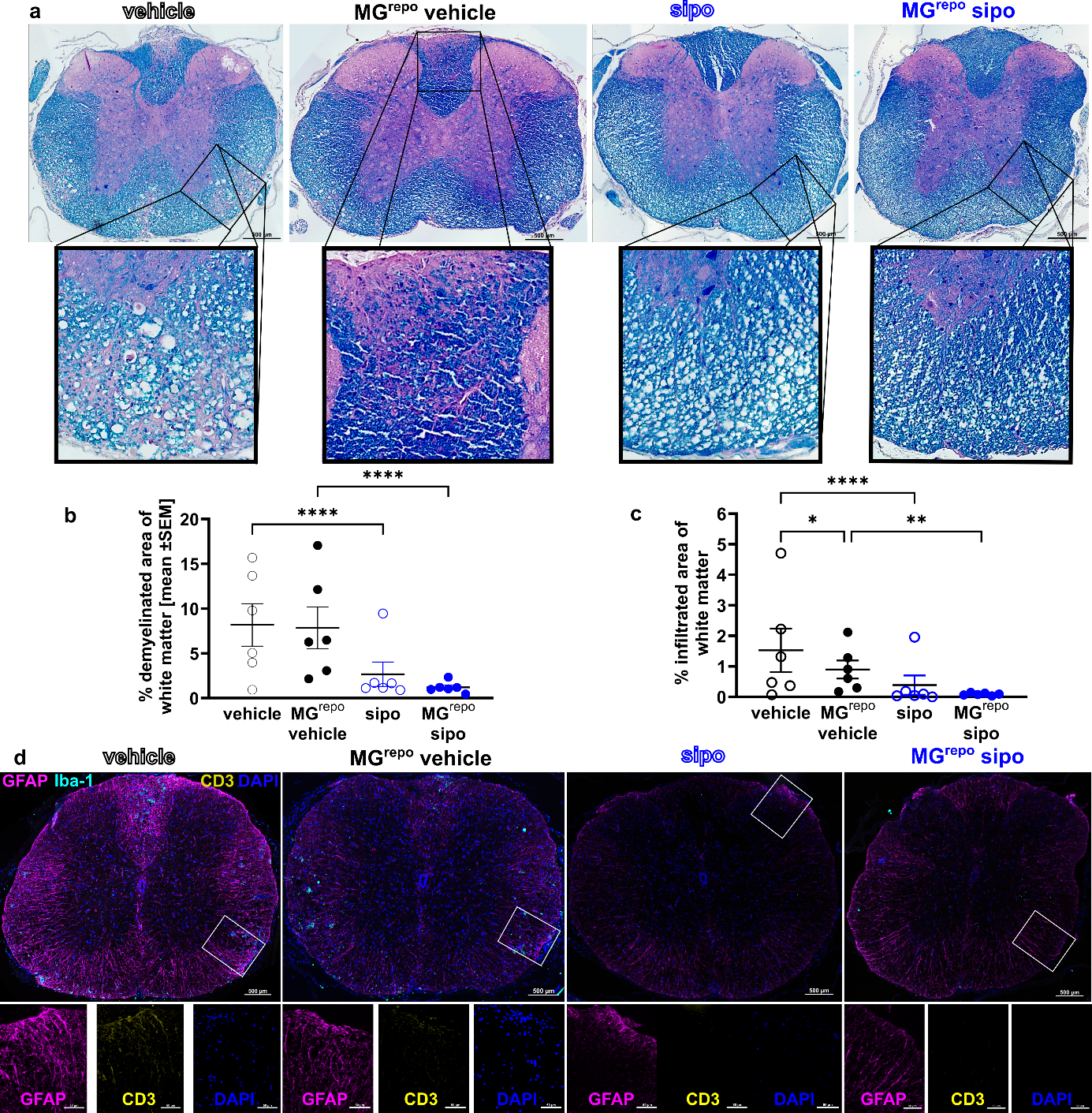

**Fig. S5. spinal cord stainings.** Spinal cord paraffin-embedded cross-sections of experimental autoimmune encephalomyelitis (EAE) mice with daily 1 µg siponimod (sipo) treatment and day 28 to 30 diphtheria toxin injection for microglia depletion and repopulation at day 35 after immunization (MG^repo^) for EAE. (a) exemplary stainings of hematoxylin and eosin (H&E) with luxol fast blue (LFB). (b) demyelinated area of the white matter evaluated from LFB and (c) infiltrated area of the white matter analyzed from H&E staining. (d) exemplary pictures of immunostaining with GFAP, Iba-1 and CD3 to detect immune cell composition in the infiltrates. Nuclei were stained with DAPI. MG – microglia, Data are shown as mean ±SEM. Statistical significance was tested with 2-way ANOVA with Šídák's multiple comparisons test with mean, SEM and n per animal with n=6 animals per group. Scale bars are 500 µm in the overview pictures and 50 µm in the detail pictures. P-values are depicted as * - p≤0.05, ** - p≤0.01, *** - p≤0.001, **** - p≤0.0001.

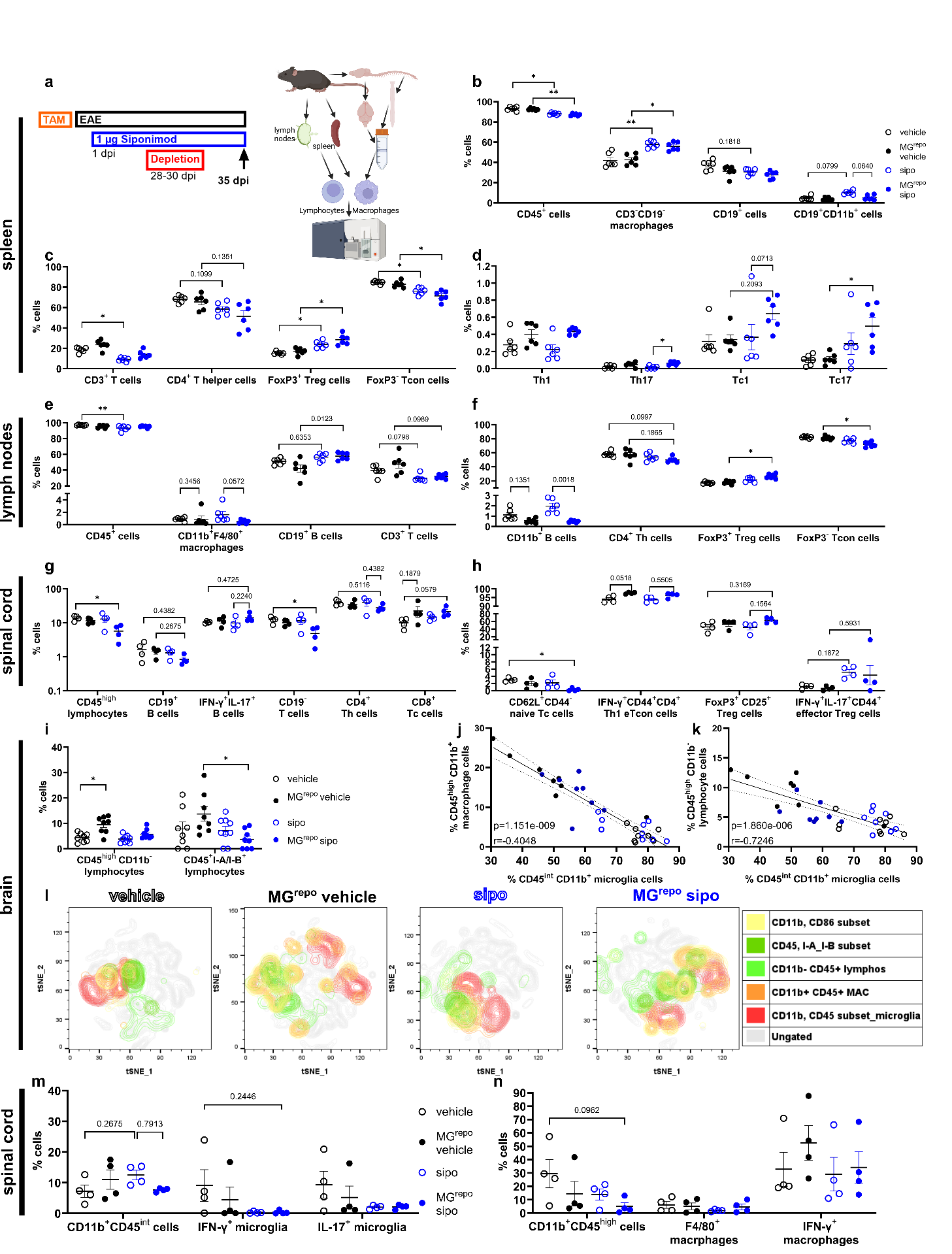
**Fig. S6. Lymphocyte and macrophage analyses after preventive siponimod treatment and microglia repopulation in chronic EAE.** Experimental design (a). Flow cytometry analysis of immune cell subsets in spleen (b-d, n=6), lymph nodes (e-f, n=6), spinal cord (g,h, n=4) and brain (i-l, n=8-9) including unsupervised clustering of the living cells with the samples per group in one concatenated file based on the gating presented in the gating strategy (i,l) outlining the efficacy of low dose siponimod (sipo) on peripheral immune cells and brain infiltration changes induced by microglia repopulation (MG^repo^). Correlation of macrophages and lymphocytes with microglia (j,k). Flow cytometry of microglia (m) and macrophages (n) in the spinal cord (n=4). Data are shown as mean ±SEM (b-i). Data were analyzed using Kruskal-Wallis tests with Dunn’s multiple comparisons tests (b-i) and Spearman correlation (j,k). (a) was created with Biorender. P-values are depicted as * - p≤0.05, ** - p≤0.01, *** - p≤0.001, **** - p≤0.0001.

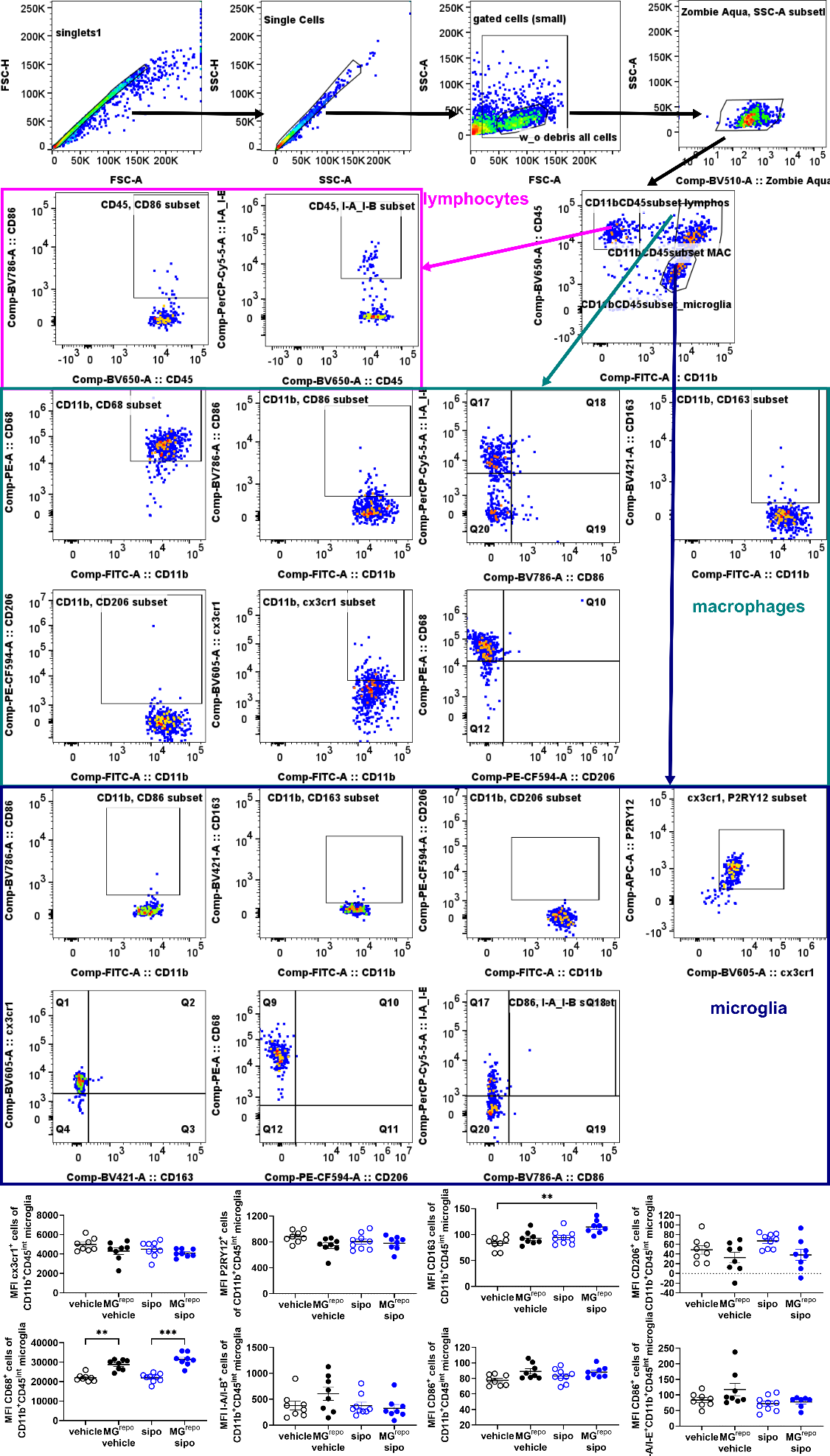

**Fig. S7: Gating strategy for brain immune cell and microglia flow cytometry with FlowJo.** Staining with microglia panel. Below the geometric mean fluorescence intensities of the analyzed parameters for the microglia population.

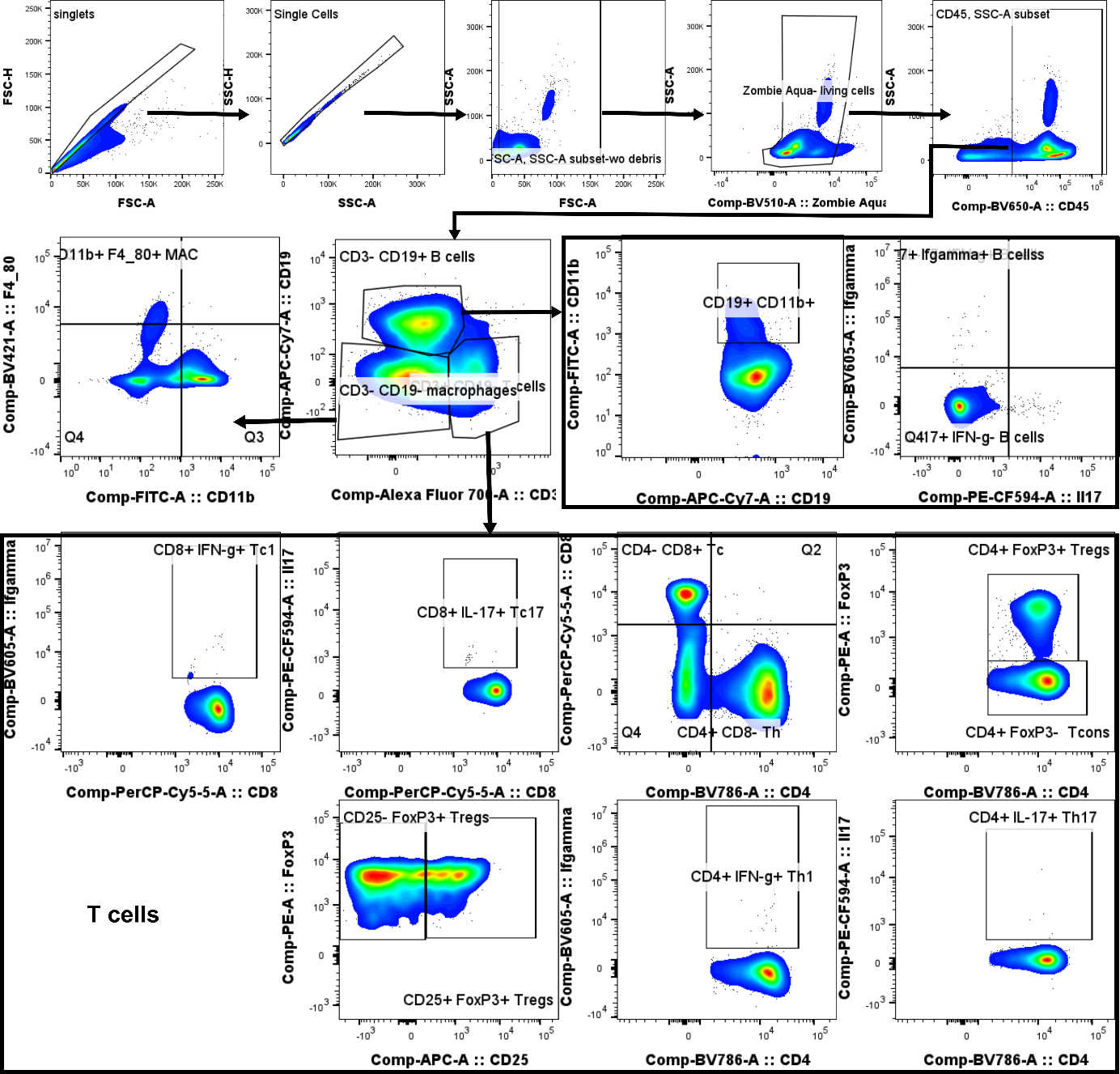

**Fig. S8. gating strategy for blood, spleen and lymph nodes flow cytometry with FlowJo.** Spleen gates are shown exemplary.

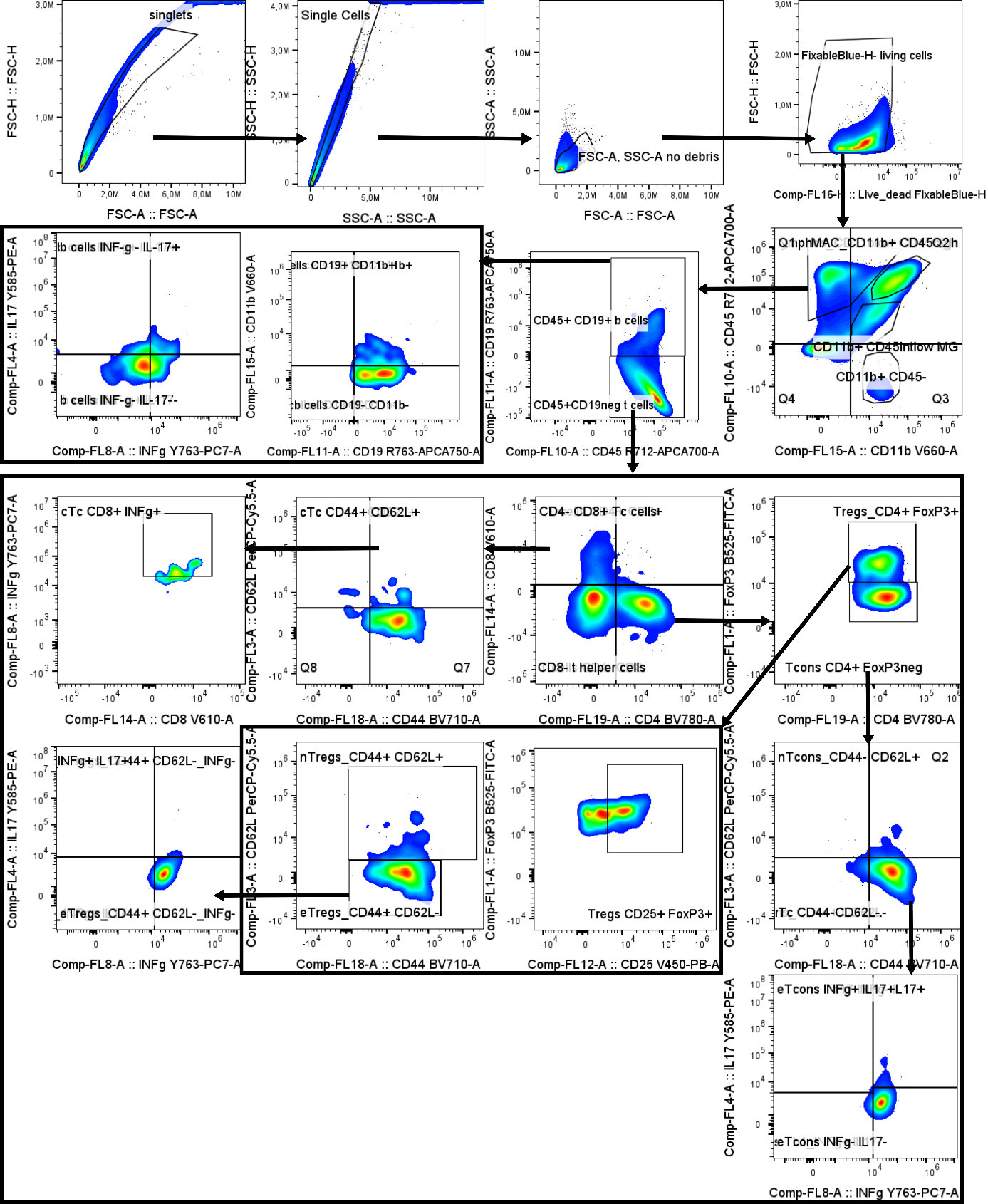

**Fig. S9. gating strategy immunophenotyping of Percoll-isolated spinal cord immune cells with FlowJo.**

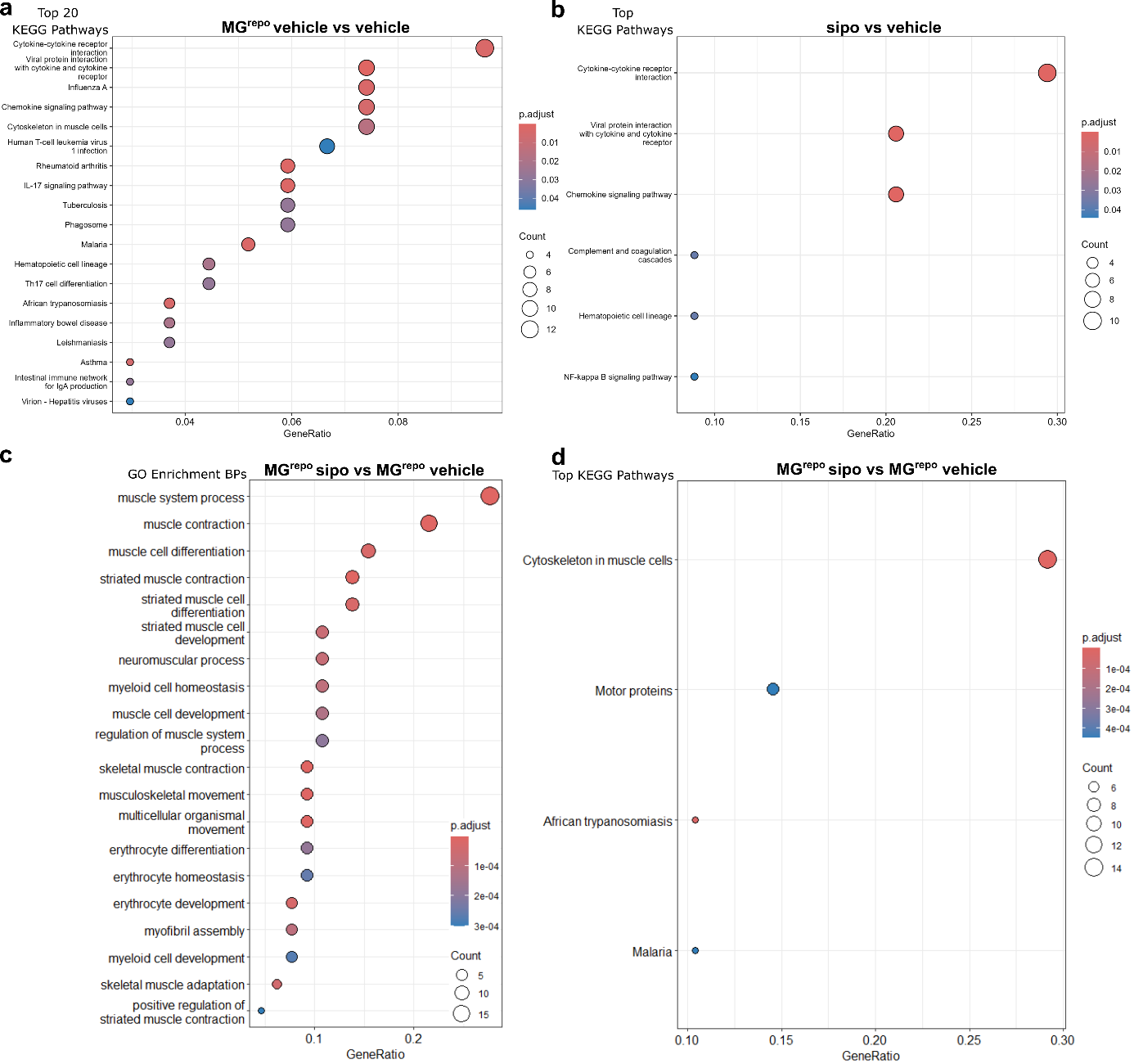

**Fig. S10. Analyses of pathways and biological processes associated with deregulated genes in the spinal cord.** Lumbar spinal cord RNA sequencing analyses of pooled EAE mice per group with daily 1 µg siponimod (sipo) treatment and day 28 to 30 diphtheria toxin injection for microglia depletion and repopulation at day 35 after immunization (MG^repo^) for EAE. The DEGs were analyzed for KEGG pathways and GO terms of biological processes.

#### Tables

Table S1. Supplemental protein quantification mass spectrometry data used to create Figure 6.

Table S2. Supplemental RNA sequencing data used to create Figure 7 and Fig. S9.

Table S6. Supplemental statistical stratification for sex differences.

**Table S3. antibodies for immunostainings.** ICC – immunocytochemistry, IHC-F – immunocytochemistry of frozen sections, IHC-P – immunocytochemistry of paraffin embedded sections, FC – flow cytometry, AF – Alexa Fluor, α – anti, rt – rat, ms – mouse, ch – chicken, rb – rabbit, dk – donkey, gt - goat.

| **target** | **clone** | **dilution** | **producer** | **reference number** |
| --- | --- | --- | --- | --- |
| CD3 | CD3-12 | 1:200 (IHC-P) | BioRad | MCA1477 |
| CD86 | GL-1 | 1:500 (ICC) | BioLegend | 105002 |
| GFAP | GA5 | 1:800 (IHC-P)  1:500 (IHC-F) | Merck | MAB360 |
| Iba‑1 | Ch311H9 | 1:400 (ICC)  1:500 (IHC-F) | Synaptic Systems | 234 009 |
| Iba‑1 | - | 1:100 (IHC-P) | Fujifilm Wako | 019-19741 |
| iNOS | CXNFT | 1:500 (ICC) | Invitrogen | 14-5920-82 |
| MBP | SMI99 | 1:500 (IHC-PF) | BioLegend | 808403 |
| NFH | SMI31 | 1:200 (IHC-P/F) | Covance | SMI-31P-100 |
| ßIII-Tubulin | - | 1:1000 (ICC) | Sigma Aldrich | T-2200-200UL |
| CD11b | M1/70 | 1:200 (FC) | BioLegend | 101206 |
| FoxP3 | FJK16S | 1:200 (FC) | Invitrogen | 12-5773-82 |
| IL-17A | TC11-18H10.1 | 1:50 (FC) | BioLegend | 506937 |
| CD8a | 53-6.7 | 1:200 (FC) | BioLegend | 100733 |
| CD25 | PC61.5 | 1:200 (FC) | eBiosciences | 17-0251-81A |
| CD3 | 17A2 | 1:100 (FC) | BioLegend | 100215 |
| CD19 | 1D3 | 1:200 (FC) | BD | 561 737 |
| F4/80 | BM8 | 1:50 (FC) | BioLegend | 123131 |
| Zombie Aqua | - | 1:1500 (FC) | BioLegend | 423101 |
| IFN-gamma | XMG1.2 | 1:100 (FC) | BioLegend | 505839 |
| CD45 | 30-F11 | 1:50 (FC) | BioLegend | 103151 |
| CD4 | RM4-5 | 1:200 (FC) | BioLegend | 100551 |
| P2RY12 | S16007D | 1:200 (FC) | BioLegend | 848006 |
| CD163 | S15049I | 1:400 (FC) | BioLegend | 155309 |
| CX3CR1 | SA011F11 | 1:50 (FC) | BioLegend | 149027 |
| CD206 | C068C2 | 1:50 (FC) | BioLegend | 171732 |
| CD68 | FA-11 | 1:100 (FC) | BioLegend | 137014 |
| I-A/I-E | M5/114.15.2 | 1:400 (FC) | BioLegend | 107626 |
| CD86 | GL-1 | 1:200 (FC) | BioLegend | 105043 |
| FoxP3 | FJK-16s | 1:200 (FC) | eBioscience | 53-5773-82 |
| CD62L | MEL-14 | 1:1000 (FC) | eBioscience | 45-0621-82 |
| IL-17A | TC11-18H10.1 | 1:50 (FC) | BioLegend | 506904 |
| IFN-gamma | XMG1.2 | 1:50 (FC) | eBioscience | 25-7311-82 |
| F4/80 | BM8 | 1:1000 (FC) | BioLegend | 123116 |
| CD45 | 30-F11 | 1:500 (FC) | BioLegend | 103128 |
| CD19 | 6D5 | 1:1000 (FC) | BioLegend | 115530 |
| CD25 | PC61 | 1:250 (FC) | BioLegend | 102043 |
| CD3 | 500A2 | 1:100 (FC) | BD | 560771 |
| CD8a | 53-6.7 | 1:1000 (FC) | BioLegend | 100744 |
| CD11b | M1/70 | 1:1000 (FC) | BioLegend | 101259 |
| CD44 | IM7 | 1:1000 (FC) | BioLegend | 103057 |
| CD4 | RM4-5 | 1:1000 (FC) | BioLegend | 100516 |
| LIVE/DEAD™ Fixable Blue Dead Cell Stain Kit | - | 1:1000 (FC) | Invitrogen | L34961 |
| AF488rtαms | - | 1:100 (IHC-P) | BioLegend | 406718 |
| AF647rtαms | - | 1:100 (IHC-P) | BioLegend | 406618 |
| Cy3dkαch | - | 1:100 (ICC)  1:400 (IHC-F) | Millipore | AP194C |
| AF488dkαrt | - | 1:500 (ICC) | Invitrogen | A48269 |
| AF555dkαrt | - | 1:1000 (IHC-P)  1:500 (IHC-F) | Invitrogen | A48270 |
| AF647gtαms | - | 1:1000 (IHC-P)  1:500 (IHC-F) | Invitrogen | A21237 |
| AF546gtαms | - | 1:1000 (IHC-P) | Invitrogen | A-11071 |
| AF488gtαrb | - | 1:1000 (IHC-P) | Invitrogen | A32731 |
| Cy5gtαrb | - | 1:400 (ICC)  1:1000 (IHC-F) | Abcam | ab97077 |
| AF488dkαrb | - | 1:500 (IHC-F) | Invitrogen | A21206 |

**Table S4. primer pairs for quantitative reverse transcriptase PCR.**

| **Gene name** | **Oligonucleotide name** | **Sequence (5'-3')** |
| --- | --- | --- |
| *Actb* | β-Actin fw. | GAC CTC TAT GCC AAC ACA GT |
|  | β-Actin rev. | AGT ACT TGC GCT CAG GAG GA |
| *Arg1* | ARG-1 fw. | GCA GAT TCC CAG AGC TGG TT |
|  | ARG-1 rev. | CTT TCT CAA AAG GAC AGC CTC G |
| *Cd86* | CD86 fw. | CTT ACG GAA GCA CCC ACG AT |
|  | CD86 rev. | TGT AAA TGG GCA CGG CAG AT |
| *Cxcl2* | CXCL-2 fw. | CAAAGGCAAGGCTAACTGACC |
|  | CXCL-2 rev. | CATCAGGTACGATCCAGGCTT |
| *Il1b* | IL-1β fw. | GCC ACC TTT TGA CAG TGA TGA G |
|  | IL-1β rev. | TGA TGT GCT GCT GCG AGA TT |
| *Mrc1* | CD206 fw. | AAC CAG TTC CTT CAG CTC GG |
|  | CD206 rev. | CTG ATT AGG GCA GCC GGT AG |
| *Tbp* | TBP fw. | AGC TCT GGA ATT GTA CCG CA |
|  | TBP rev. | TGA CTG CAG CAA ATC GCT TG |
| *Tlr4* | TLR4-fw | GGACTCTGATCATGGCACTG |
|  | TLR4-rw | CTGATCCATGCATTGGTAGGT |
| *Tnf* | TNFα fw. | ATG GCC TCC CTC TCA TCA GT |
|  | TNFα rev. | TGG TTT GCT ACG ACG TGG G |
| *Trem2* | Trem2-fw | TGGGACCTCTCCACCAGTT |
|  | Trem2-rv | GTGGTGTTGAGGGCTTGG |
| *Ym1, Chil3 Chi3l3* | Ym1 fw. | GGT CTG AAA GAC AAG AAC ACT |
|  | Ym1 rev. | GAG ACC ATG GCA CTG AAC G |

**Table S5. components of cell culture media.** Pen – penicillin, strep – streptomycin, MEM – minimal essential medium, DMEM – dulbecco’s modified eagle medium.

| **medium name** | **component** | **concentration** |
| --- | --- | --- |
| Supplemented MEM | MEM |  |
|  | GlutaMAX | 1x |
|  | pen/strep | 100 µg/mL |
|  | non-essential amino acids | 1x |
|  | sodium pyruvate | 1x |
|  | dextrose | 0.10% |
|  | fetal bovine serum | 10% |
| supplemented DMEM | DMEM/F-12, GlutaMAX™ Supplement (Gibco) |  |
|  | fetal bovine serum | 10% |
|  | pen/strep | 1% |
| complete Neurobasal | Neurobasal (Gibco) |  |
|  | B27 | 1x |
|  | pen/strep | 1% |
|  | GlutaMAX | 1x |

##### Supplement References

1.	*Nf-Core/rnaseq: Nf-Core/rnaseq V3.19.0 - Tungsten Turtle.* Zenodo; 2025.

2.	Posit team. RStudio: Integrated Development Environment for R. http://www.posit.co/

3.	Kolde R. Pheatmap: Pretty Heatmaps. R Package Version 1.0.12. https://CRAN.R-project.org/package=pheatmap

4.	Love MI, et al. Moderated estimation of fold change and dispersion for RNA-seq data with DESeq2. *Genome Biol*. 2014;15(12):550. doi:10.1186/s13059-014-0550-8

5.	Wickham H, ed. *Ggplot2: Elegant Graphics for Data Analysis*. Springer International Publishing; 2016.

6.	Tarazona S, et al. Data quality aware analysis of differential expression in RNA-seq with NOISeq R/Bioc package. *Nucleic Acids Res*. 2015;43(21):e140. doi:10.1093/nar/gkv711

7.	Tarazona S, et al. Differential expression in RNA-seq: a matter of depth. *Genome Res*. 2011;21(12):2213-2223. doi:10.1101/gr.124321.111

8.	Yu G, et al. clusterProfiler: an R package for comparing biological themes among gene clusters. *OMICS*. 2012;16(5):284-287. doi:10.1089/omi.2011.0118

9.	Bioconductor. BiomaRt Package. https://bioconductor.org/packages/release/bioc/html/biomaRt.html

10.	Bioconductor. ClusterProfiler Package. https://bioconductor.org/packages/release/bioc/html/clusterProfiler.html

11.	*Org.Mm.Eg.Db: Genome Wide Annotation for Mouse.* Bioconductor; 2023.

12.	R Core Team. Graphics: The R Graphics Package: Version 3.6.2. https://www.rdocumentation.org/packages/graphics/versions/3.6.2/topics/dotchart

13.	Sielaff M, et al. Evaluation of FASP, SP3, and iST Protocols for Proteomic Sample Preparation in the Low Microgram Range. *J Proteome Res*. 2017;16(11):4060-4072. doi:10.1021/acs.jproteome.7b00433

14.	Tyanova S, et al. The Perseus computational platform for comprehensive analysis of (prote)omics data. *Nat Methods*. 2016;13(9):731-740. doi:10.1038/nmeth.3901

15.	Wulf M, et al. Neuromelanin granules of the substantia nigra: proteomic profile provides links to tyrosine hydroxylase, stress granules and lysosomes. *J Neural Transm (Vienna)*. 2022;129(10):1257-1270. doi:10.1007/s00702-022-02530-4
